# Unique unfoldase/aggregase activity of a molecular chaperone that dysregulates a translational elongation factor

**DOI:** 10.1101/492876

**Authors:** Ku-Sung Jo, Ji-Hun Kim, Kyoung-Seok Ryu, Young-Ho Lee, Che-Yeon Wang, Joo-Seong Kang, Yoo-Sup Lee, Min-Duk Seo, Hyung-Sik Won

## Abstract

The various chaperone activities of heat shock proteins contribute to ensuring cellular proteostasis. Here, we demonstrate the non-canonical unfoldase activity as an inherent functionality of the prokaryotic molecular chaperone Hsp33. The holding-inactive, reduced form of Hsp33 (^R^Hsp33) strongly bound to the translational elongation factor, EF-Tu, and catalyzed the EF-Tu aggregation via evoking its aberrant folding, resulting in its susceptibility to proteolytic degradation by Lon. This interaction was critically mediated by the redox-switch domain of ^R^Hsp33 and the guanine nucleotide-binding domain of EF-Tu. The ^R^Hsp33-induced *in vivo* aggregation of EF-Tu upon heat shock was evident in a Lon-deficient strain and inhibited cell growth. Unlike wild-type *Escherichia coli*, the strain lacking both Hsp33 and Lon showed a non-reduced level of EF-Tu and diminished capability of counteracting heat shock. These findings suggest that the unique unfoldase/aggregase activity of Hsp33 potentially involved in protein turnover confers a cellular survival advantage under heat-stressed conditions.

## Introduction

Molecular chaperones such as heat shock proteins (Hsps) are widely recognized as vital mediators of cellular protein conformation and turnover (i.e., the balance between protein synthesis and protein degradation) (Craig *et al*, 1994). Owing to the flexible and labile nature of protein structures, a complex and dynamic network of molecular chaperones constituting an elaborate machinery of protein quality control (PQC) protects the cellular proteome under both normal and stressed conditions (Balchin *et al*, 2016; Bershtein *et al*, 2013; Dahl *et al*, 2015). In particular, as misfolded and/or denatured proteins are prone to aberrant behavior such as by forming irreversible aggregates that are deleterious to cells, molecular chaperones assist in the correct *de novo* folding and refolding of proteins, and function to prevent proteins from aberrant folding and aggregation (Doyle *et al*, 2013; Hartl *et al*, 2011). In some cases, molecular chaperones are also associated with the timely removal of irreversibly misfolded proteins. As such, molecular chaperones play central roles in ensuring cellular proteostasis (i.e., proteome homeostasis) that also requires a regulated process of protein turnover (Klaip *et al*, 2017; Taipale *et al*, 2014; Visscher *et al*, 2016). Therefore, various molecular activities are associated with chaperone functions, including foldase, refoldase, unfoldase, holding, disaggregase, translocase, and targetase activities (Mattoo & Goloubinoff, 2014). Among these, the unfoldase activity of molecular chaperones, which is somewhat in conflict with its overall function in conformational maintenance, catalyzes the unfolding of stable misfolded proteins to convert them into natively refoldable species. This unfoldase activity is therefore generally linked to other chaperone activities such as in the sequential reaction for disaggregation-unfolding-refolding of stable soluble aggregates (Mattoo *et al,* 2014; Priya *et al*, 2013), or in the case of AAA+ proteases that unfold misfolded substrate proteins for subsequent degradation (Van Melderen & Aertsen, 2009; Vieux *et al*, 2013). In both cases, most unfoldase reactions are ATP-consuming processes. Here, we report a novel type of unfoldase activity identified in the prokaryotic molecular chaperone Hsp33, which is an ATP-independent process inducing irreversible aggregation of the substrate protein.

Hsp33 was originally discovered as a heat-inducible, but post-translationally activated chaperone (Dahl *et al*, 2015; Jakob *et al*, 1999). The protein requires dual stressors of heat and oxidation (Ilbert *et al*, 2007), or a severe acute oxidative stress signal (Winter *et al*, 2008) to display its ATP-independent holding activity, which results in binding to the unfolding intermediates of client proteins to prevent their ultimate, irreversible denaturation. The reduced form of Hsp33 (^R^Hsp33; refer to *figure supplement 1A* for a structural depiction), which is inactive in terms of the holding chaperone function, has a unique fold of the redox-switch domain (RSD; residues 232–294) that binds a zinc ion via four conserved cysteines (C232, C234, C265, and C268) (Won *et al*, 2004). Thus, under oxidative heat or kinetically fast oxidative conditions, the functionally active, oxidized form of Hsp33 (^O^Hsp33) is formed via releasing zinc due to disulfide bond formation (C232-C234 and C265-C268 linkages) of the cysteines. In this holding-active conformation, the middle linker domain (MLD; residues 179–231) as well as the RSD become disordered to provide client-binding sites (Groitl *et al*, 2016; Reichmann *et al*, 2012). However, mild oxidation of ^R^Hsp33 at a non-elevated temperature predominantly produces a half-oxidized form of the protein (^hO^Hsp33) with only one disulfide bond (C265-C268) in which the RSD is unfolded, but the MLD remains folded, and the protein shows no or only slight activity (Ilbert *et al*, 2007; Lee *et al*, 2012). Subsequent studies revealed that unfolding of the RSD is not the structural determinant for functional activation of Hsp33, although it serves as a redox-sensing module (Cremers *et al*, 2010; Lee *et al*, 2015). This observation, together with the facts that Hsp33 is expressed at a basal level even under non-stressed conditions (Dahl *et al*, 2015) and heat shock itself does not provide an ample stimulus for the thermally overexpressed Hsp33 to achieve its holding chaperone activity, led us to speculate that ^R^Hsp33 may possess its own specific functionality via the unique fold of the RSD. In this regard, we considered the findings of controversial reports (Wholey & Jakob, 2012; Bruel *et al*, 2012; Voth & Jakob, 2017) investigating the effects of Hsp33 on the elongation factor thermos-unstable (EF-Tu) protein, a translational GTPase playing an essential role in translation elongation by delivering aminoacyl-tRNA to the ribosomal A site (Maracci & Rodnina, 2016; Noble & Song, 2008). For example, Wholey and Jakob observed that Hsp33 protected EF-Tu against oxidative degradation in *Vibrio cholera* (Wholey & Jakob, 2012), whereas Bruel et al. demonstrated that Hsp33 overexpression targeted EF-Tu for degradation in an *Escherichia coli* strain lacking trigger factor (TF) and DnaK (Bruel *et al*, 2012). However, the molecular interaction of Hsp33 with EF-Tu has not yet been investigated at the molecular structural level.

In this context, the aim of the present study was to uncover the novel functionality of Hsp33 and to underpin its structural basis. In addition to verifying the direct molecular interaction, we aimed to identify which of the multiple conformations of Hsp33 (^R^Hsp33, ^hO^Hsp33, and ^O^Hsp33) is primarily responsible for the EF-Tu interaction, to determine whether it protects or destabilizes EF-Tu, and to identify the structural and functional consequences of the interaction. Overall, the results demonstrate that ^R^Hsp33 specifically displays unique unfoldase/aggregase activity against EF-Tu, and this molecular action in cells, coupled with Lon protease, confers an obvious survival advantage under heat shock conditions.

## Results

### Oligomerizing and aggregating tendency of EF-Tu

The guanine nucleotide-binding domain (G-domain) of EF-Tu in cells binds a functional magnesium ion, which is then transiently dissociated for guanine nucleotide exchange of the protein (Maracci & Rodnina, 2016; Noble & Song, 2008). Our initial preparation of recombinant EF-Tu using no particular additives resulted in a heterogeneous mixture of the protein, indicative of various oligomeric states (red trace in Fig 1A). Specifically, the gel-filtration profile of the purified EF-Tu (theoretically 46 kDa including the hexahistidine tag) showed three distinct eluates of the monomer (the last eluate with an estimated hydrodynamic size of 45 kDa), dimer (middle, 81 kDa), and high-order oligomer (column void-volume eluate) species, respectively. The particle sizes deduced for the first two eluates by dynamic light scattering also supported the presence of a dimeric (81 kDa with a 4-nm radius) and 20-mer oligomeric (883 kDa with an 11-nm radius) species (Fig EV2). In addition, the purified EF-Tu showed a potent tendency of time-dependent aggregation leading to gradual precipitation during a few days of storage at room temperature. Storage at a low temperature (4°C) for more than 4 days also resulted in nearly complete conversion to the oligomeric state, thereby showing almost a single eluate at the column void volume (blue trace in Fig 1A).

**Figure 1.**
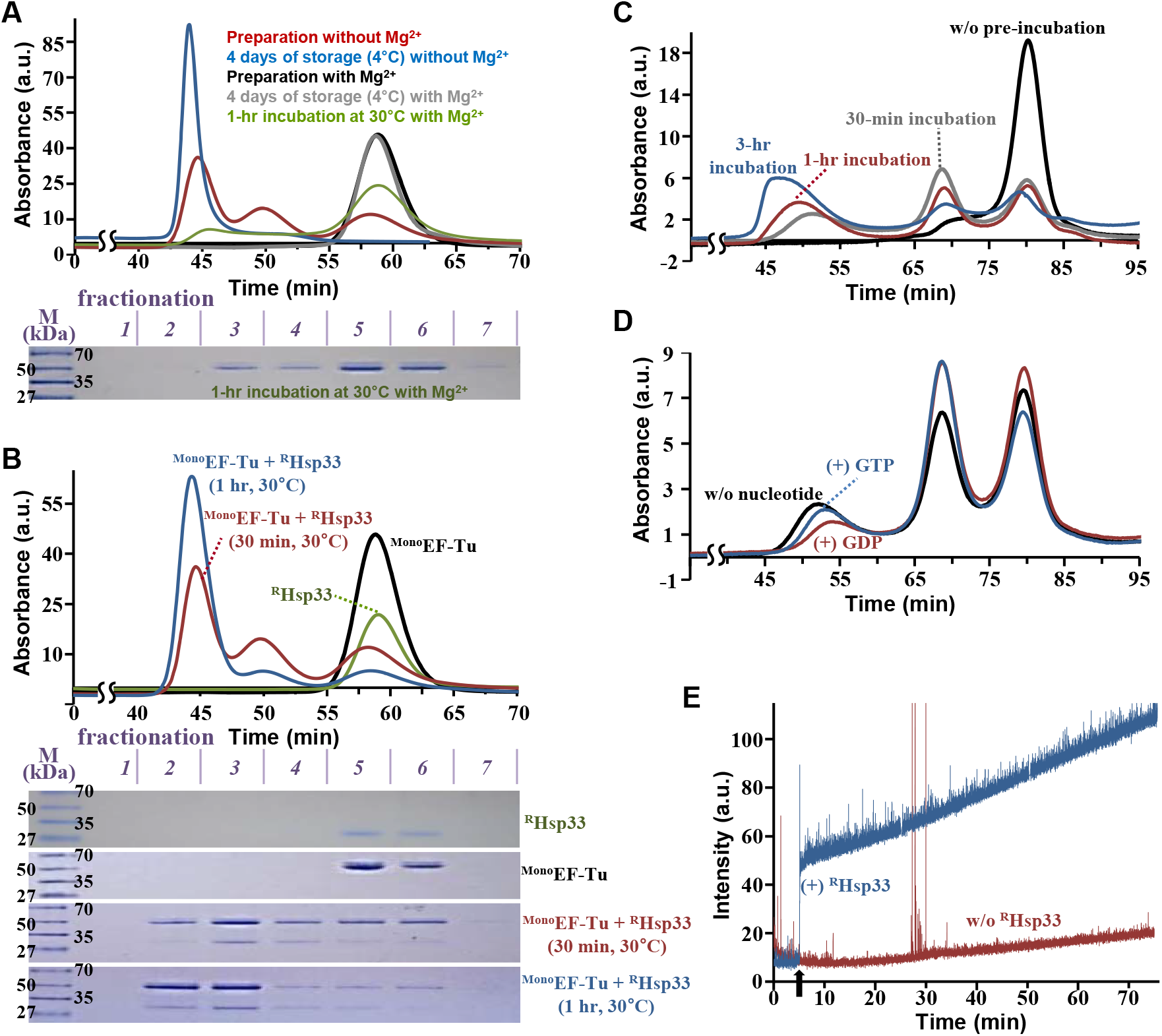
Oligomerization of EF-Tu. The analytical gel-filtration assay was performed at room temperature using a Superdex 75 (**A** and **B**) or Superdex 200 (**C** and **D**) column, whereas light scattering was monitored at 30°C (**E**). **A.** EF-Tu (50 μM) solutions prepared in the absence (red and blue lines) and presence (black, gray, and green lines) of Mg^2+^ were analyzed instantly after purification (red and black lines), after four days of storage at 4°C (blue and gray lines) and after 1-h incubation at 30°C (green line), respectively. SDS-PAGE (12% Tricine gel) images for individual fractions (collected every 5 min) of the green profile are shown in the bottom panel. **B.** Standard profiles for isolated ^R^Hsp33 and ^Mono^EF-Tu are presented by green and black traces, respectively. The mixture of ^Mono^EF-Tu (50 μM) and ^R^Hsp33 (25 μM) was incubated at 30°C for 30 min (red) or for 1 h (blue) prior to injection. All individual eluents were fractionated every 5 min and resolved by 12% Tricine-SDS-PAGE (lower panel; M, molecular size marker). **C.** The ^Mono^EF-Tu (15 μM) and ^R^Hsp33 (15 μM) mixture was injected immediately after mixing (black) and after incubation at 30°C for 30 min (gray), 1 h (red), or 3 h (blue). **D.** The ^Mono^EF-Tu (15 μM) and ^R^Hsp33 (15 μM) mixture without nucleotides (black) or containing 60 μM of GTP (blue) or GDP (red) migrated after 1 h pre-incubation at room temperature (approximately 22°C). **E.** Light (400 nm) scattering of ^Mono^EF-Tu (100 μM) solution in the absence (red) and presence (blue) of ^R^Hsp33 (50 μM; added at 5 min of incubation, as indicated by a black arrow) was monitored over incubation time.

In contrast, the gel-filtration result for a different preparation with continuous use of excess Mg^2+^ during all steps of protein expression and purification showed a single species of the elution corresponding to its monomeric size (black trace in Fig 1A), which was maintained during storage at 4°C for more than 4 days (gray trace in Fig 1A). However, adding ethylenediaminetetraacetic acid (EDTA) to the Mg^2+^-containing monomeric EF-Tu (^Mono^EF-Tu) solution resulted in severe precipitation (data not shown), which was likely due to rapid oligomerization of the protein. These results indicated that the Mg^2+^ ion bound to EF-Tu is crucial for the stability of the protein. In addition, the oligomerization of Mg^2+^-bound EF-Tu also readily proceeded at elevated temperatures (green trace in Fig 1A) and was accelerated at higher temperatures, indicating its intrinsically thermo-unstable property.

### Reduced form-specific binding of Hsp33 to EF-Tu

A pull-down assay was performed to examine the Hsp33 binding to EF-Tu by employing the hexahistidine-tagged ^Mono^EF-Tu as bait for binding to the Ni^2+^-affinity resin, along with three Hsp33 preparations at different redox statuses (^R^Hsp33, ^hO^Hsp33, and ^O^Hsp33) as prey, respectively. ^R^Hsp33 snapped at the resin-bound EF-Tu, whereas ^hO^Hsp33 showed no significant binding (Fig 2A). As anticipated, ^O^Hsp33 also did not bind to EF-Tu (data not shown). EF-Tu binding to ^R^Hsp33 was then examined by nuclear magnetic resonance (NMR) spectroscopy (Fig 2B). The [^1^H/^15^N]TROSY spectrum of ^R^Hsp33 showed significant line broadening upon the addition of ^Mono^EF-Tu (*Figure 2B*), indicating formation of a complex with the protein. Subsequently, we attempted to measure the binding affinity using isothermal titration calorimetry (ITC). Consistent with the pull-down assay results, no significant binding of ^hO^Hsp33 and ^O^Hsp33 to ^Mono^EF-Tu was observed (Fig EV3). Unexpectedly, the ITC thermogram for ^R^Hsp33 binding to ^Mono^EF-Tu showed an unusual trace characterized by continuous endothermic reactions following exothermic pulses (Fig 2C). This abnormal thermogram was ultimately explained by the conformational change of EF-Tu after ^R^Hsp33 binding (see Discussion). However, the exothermic traces with endothermic interference inevitably hindered the ability to conduct a reliable analysis to estimate thermodynamic parameters. Alternatively, we measured the ^R^Hsp33 binding to the preformed oligomeric EF-Tu (^Oligo^EF-Tu; void-volume fraction as shown in *Figure 1A*), which permitted a well-fitted estimation (Fig 2C): *K*_d_ of 0.58 ± 0.13 μM (Δ*H* = −78.6 ± 3.47 kJ·mol^-1^; Δ*G* = −35.7 kJ·mol^-1^) with a stoichiometry of approximately two (N = 2.37 ± 0.06) ^R^Hsp33 molecules to one EF-Tu oligomer, using the aforementioned 20-mer oligomeric size for EF-Tu. Thus, the chaperone-inactive, reduced form of Hsp33, but not oxidized forms, was revealed to be relevant to both ^Mono^EF-Tu and ^Oligo^EF-Tu binding.

**Figure 2.**
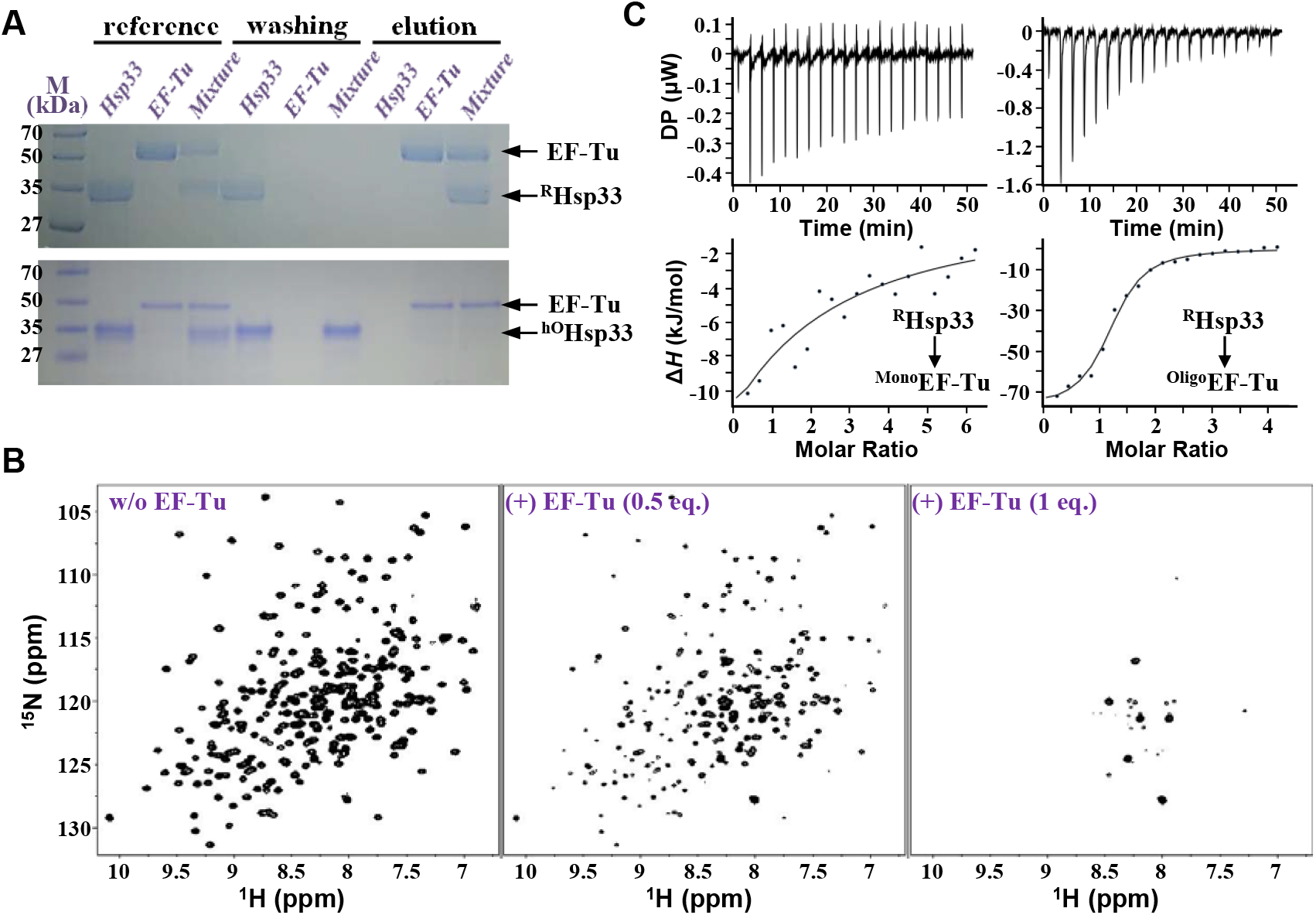
Molecular interaction between ^R^Hsp33 and EF-Tu. **A.** Pull-down assay using His-tag-fused ^Mono^EF-Tu as a bait that binds to the resin, while employing ^R^Hsp33 (upper panel) or ^hO^Hsp33 (lower panel) as a prey molecule. The protein solution was mixed with the resin, followed by washing and subsequent elution. Each solution was resolved by SDS-PAGE (lane M, size marker). **B.** NMR ([^1^H/^15^N]TROSY) spectrum of [^15^N]^R^Hsp33 (0.3 mM) in the absence (left) and presence of 0.5 (middle) and 1 (right) equimolar EF-Tu. **C.** ITC measurement for RHsp33 binding to ^Mono^EF-Tu (left) and ^Oligo^EF-Tu (right). Each point in the binding isotherm (lower panels) represents the integrated heat of the associated peak in the thermogram (upper panels).

### EF-Tu oligomerization and aggregation promoted by ^R^Hsp33

Considering our NMR instrumental conditions, the quite severe line broadening of the ^R^Hsp33 NMR spectrum in the presence of ^mono^EF-Tu (Fig 2B) was unusual and alluded to massive complex formation quite beyond a one-to-one binding of the two proteins. Thus, considering the intrinsic propensity of EF-Tu to oligomerization (Fig 1A), we reasoned that a huge complex system of the two proteins might be created after the binding, which was verified by gel-filtration analysis.

When the ^Mono^EF-Tu solution was incubated with ^R^Hsp33 at 30°C for 30 min, the gel-filtration profile of the mixture showed three distinct elution peaks commonly containing both proteins (see red trace and the gel electrophoresis image in Fig 1B). The last elution was attributable to small fractions of non-complexed proteins (note that the hydrodynamic size of ^R^Hsp33 is comparable with that of ^Mono^EF-Tu; see green and black traces in Fig 1B, respectively). The middle elution can be accounted for by the one-to-one complex of the proteins based on the estimated hydrodynamic size of approximately 80 kDa. The largest elution at the column void volume indicated the presence of high-order oligomers of the complex. Longer (1 h) incubation led to appreciable enlargement of the oligomer fraction with a concomitant decrement of monomeric and heterodimeric species (blue trace in *Figure 1B*). Furthermore, with longer incubation of the complex, the oligomer size became increasingly enlarged, as reflected by the shortened retention times of oligomer fractions for longer-incubated samples (Fig 1C). This continuously expanding size implied that the oligomerizing process could eventually result in irreversible aggregation.

Although it was not clear whether EF-Tu, ^R^Hsp33, or both are responsible, the observed oligomerization/aggregation was more vigorous with a higher concentration of EF-Tu (e.g., compare the relative proportion of the oligomer fraction in blue trace of *Figure 1B*, performed with 50 μM EF-Tu, with that in red trace of Fig 1C, performed with 15 μM EF-Tu), as well as at a higher incubation temperature (e.g., compare red trace of *Figure 1C*, performed at 30°C, with red trace of Fig 1D, performed at approximately 22°C). Therefore, collectively considering the heat-dependent intrinsic oligomerization of EF-Tu (green trace in Fig 1A) and the dominant content of EF-Tu in the ^R^Hsp33:EF-Tu oligomeric fractions (gel electrophoresis images in Fig 1B), we assumed that the ^R^Hsp33 binding promoted the rapid oligomerization of EF-Tu. This assumption was subsequently confirmed by observing the selective destabilization of EF-Tu in the ^R^Hsp33:EF-Tu complex (Figs 3 and 4).

**Figure 3.**
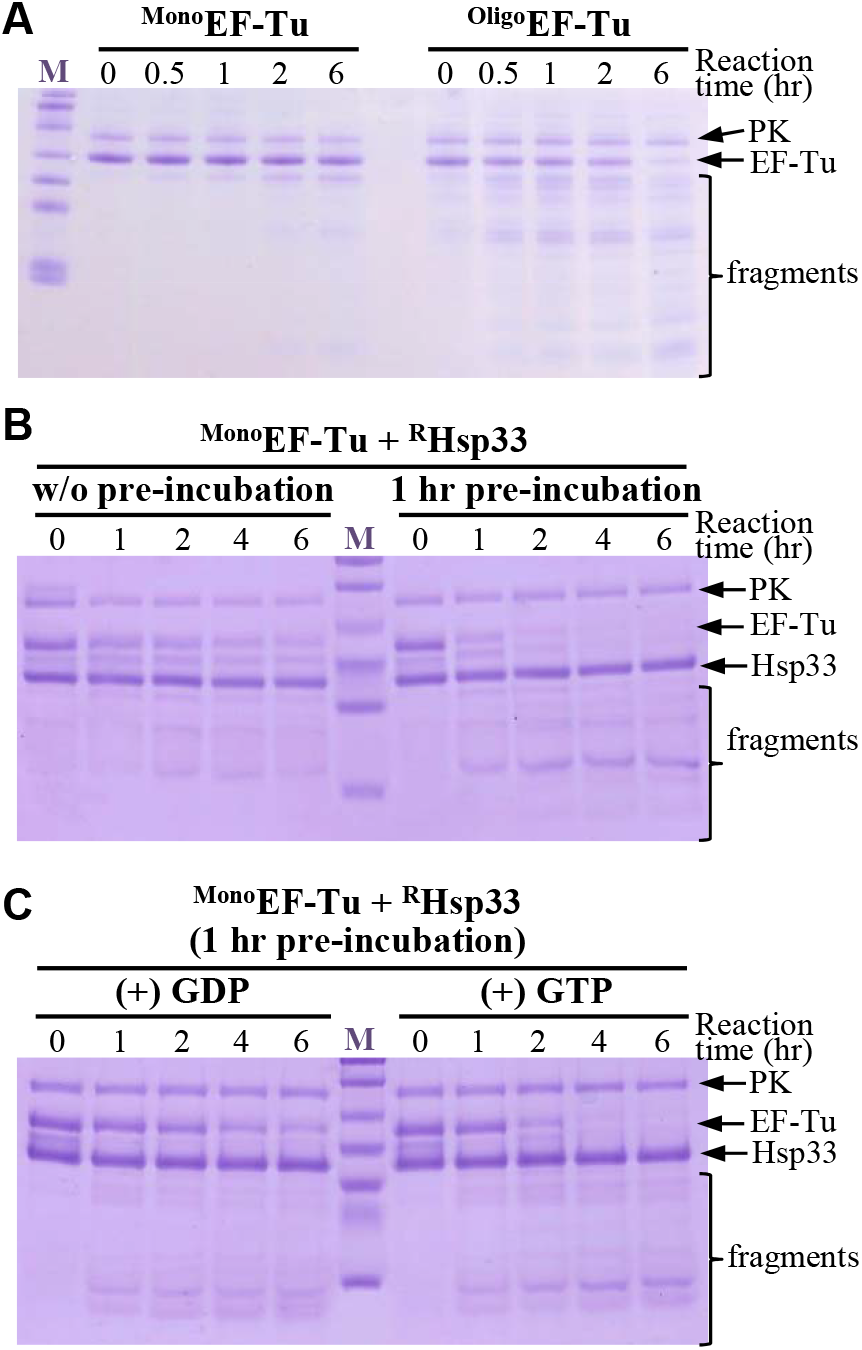
Specific digestion of oligomeric EF-Tu by Lon protease EF-Tu (10 μM) was reacted at 30°C with 0.26 nM Lon protease supplemented with pyruvate kinase (PK; 5 μM) for ATP generation, and was resolved by SDS-PAGE (12% Tricine gel). The 0-h sample was taken immediately after adding the protease to the reaction mixture. **A.** ^Mono^EF-Tu and ^Oligo^EF-Tu were subjected as substrates, respectively. **B.** The mixture of ^Mono^EF-Tu and ^R^Hsp33 (10 μM) was reacted with Lon, without or after 1-h pre-incubation. **C.** The ^Mono^EF-Tu and ^R^Hsp33 mixture containing 40 μM of GTP or GDP was pre-incubated for 1 h, followed by reaction with Lon.

**Figure 4.**
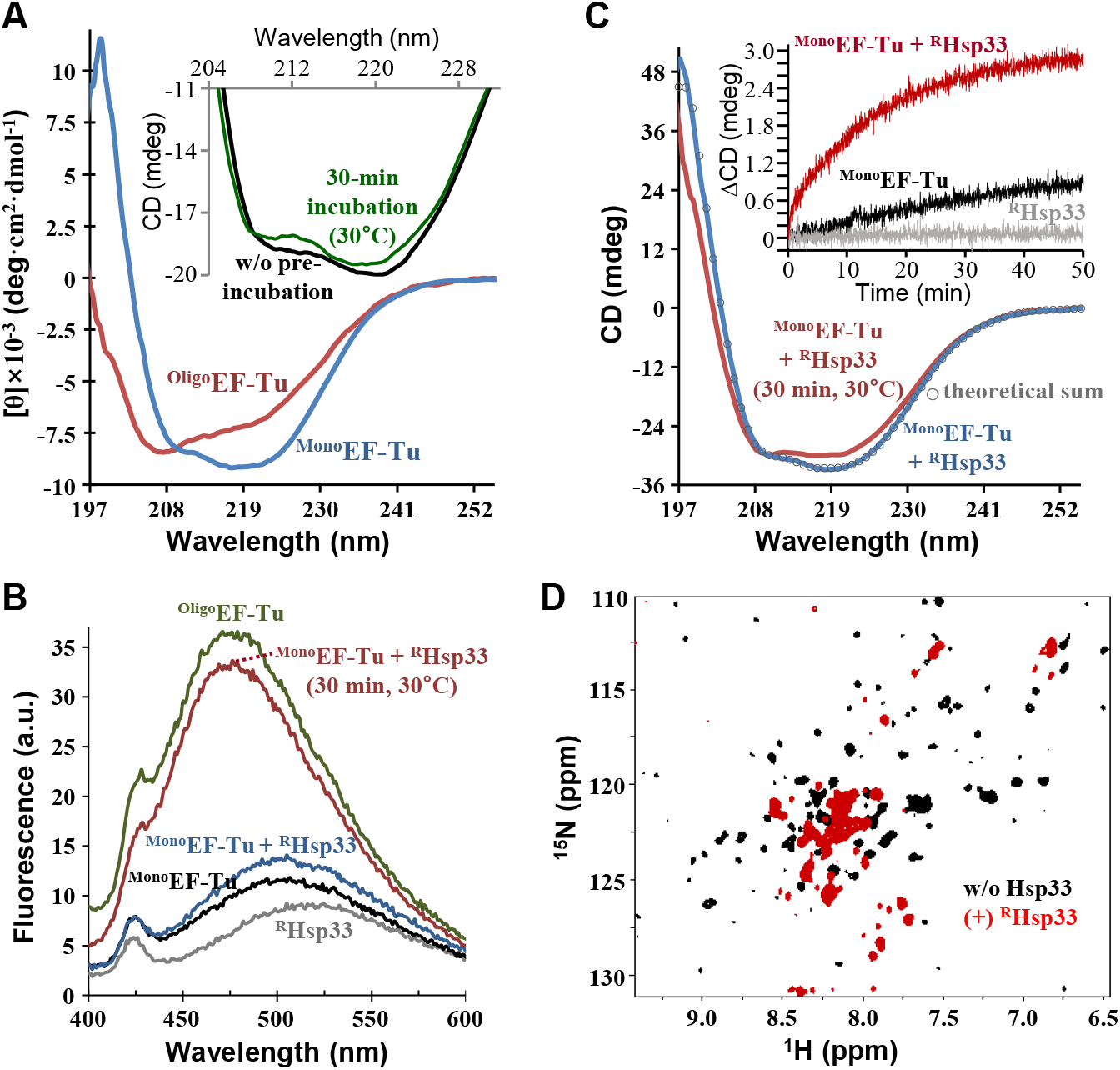
Conformational change of EF-Tu concomitant to oligomerization. **A.** Normalized ([θ], mean residue molar ellipticity) far-UV CD spectra for 8 μM of ^Mono^EF-Tu (blue) and ^Oligo^EF-Tu (red). The inset shows an enlarged region of the CD spectra for 5 μM ^Mono^EF-Tu before (black) and after (green) incubation at 30°C for 30 min. **B.** ANS fluorescence spectra of ^Mono^EF-Tu (black), ^Oligo^EF-Tu (green), ^R^Hsp33 (gray), and an equimolar (5 μM) mixture of ^Mono^EF-Tu and ^R^Hsp33 (blue, before incubation; red, after 30 min incubation at 30°C). **C.** Far-UV CD spectra of an equimolar (5 μM) mixture of ^Mono^EF-Tu and ^R^Hsp33 immediately after mixing (blue) and after 30°C incubation for 30 min (red). Open circles depict the theoretical sums of individual spectra of ^Mono^EF-Tu and ^R^Hsp33. The inset shows time-dependent CD signal changes (ΔCD) at 220 nm of 15 μM ^R^Hsp33 (gray), ^Mono^EF-Tu (black), and their mixture (red). **D.** NMR ([^1^H/^15^N]TROSY) spectra of [^15^N]EF-Tu (0.3 mM) in the absence (black) and presence (red) of equimolar ^R^Hsp33.

We also checked whether the ^R^Hsp33-mediated oligomerization of EF-Tu is affected by guanine nucleotides (GDP and GTP), whose binding is known to enhance the EF-Tu stability (Šanderová *et al,* 2004). As expected from the higher affinity of GDP than GTP (Šanderová *et al*, 2004; Thirup *et al*, 2015), the oligomerizing efficiency (relative proportion of the oligomer fraction and the oligomer size) of EF-Tu was lowest in its GDP-bound state, whereas the nucleotide-free form showed the fastest oligomerization (Fig 1D). However, none of the nucleotides inhibited Hsp33-mediated EF-Tu oligomerization, demonstrating the outperforming negative influence of ^R^Hsp33 on EF-Tu stability.

Finally, the EF-Tu aggregation was monitored over time by light scattering (Fig 1E). The intrinsic production of ^Oligo^EF-Tu in the absence of ^R^Hsp33 (green trace in Fig 1A) was reflected by the gradual, albeit slight, increase of light scattering, particularly after 20 min of ^Mono^EF-Tu incubation at 30°C (red trace in Fig 1E). The addition of ^R^Hsp33 promptly accelerated the increase of light scattering (blue trace in Fig 1E), in support of the vigorous aggregation of EF-Tu.

### Oligomer-specific degradation of EF-Tu by Lon protease

Since EF-Tu has been suggested to be a substrate of Lon protease in cells (Bruel *et al*, 2012), the susceptibility of ^Mono^EF-Tu and its intrinsically converted form ^Oligo^EF-Tu to Lon was compared (Fig 3A). Notably, ^Mono^EF-Tu appeared to be resistant to the proteolysis by Lon, whereas ^Oligo^EF-Tu was efficiently digested. The slight degradation of ^Mono^EF-Tu detected after 2-h incubation with Lon was attributable to the small fraction of oligomers created spontaneously during incubation (see green trace in Fig 1A). EF-Tu degradation by Lon was also prominent in the presence of ^R^Hsp33 (Fig 3B), implying that the ^R^Hsp33-induced ^Oligo^EF-Tu was competent for rapid digestion by Lon. Furthermore, EF-Tu degradation was highly accelerated when its mixture with ^R^Hsp33 was pre-incubated for efficient oligomerization prior to reaction with Lon (Fig 3B). Moreover, guanosine nucleotides that slightly suppressed the oligomerization (Fig 1D) also resulted in marginal retardation of the degradation (Fig 3C). Together, these results suggest that ^Oligo^EF-Tu was specifically subjected to Lon proteolysis and the Hsp33 binding could facilitate the Lon-mediated EF-Tu degradation by promoting its oligomerization. We further confirmed that Hsp33 could also be susceptible to Lon depending on its unfolding status; i.e., the isolated ^hO^Hsp33 with partial unfolding and ^O^Hsp33 with extended unfolding (Ilbert *et al*, 2007; Lee *et al*, 2015) invoked moderate and rapid digestion, respectively (Fig EV4). However, ^R^Hsp33, as well as the pyruvate kinase that was contained in the reaction mixture for ATP generation, remained intact during EF-Tu degradation (Figs 3B and 3C). Hence, a certain conformational event specific to EF-Tu could be anticipated in the ^R^Hsp33:EF-Tu complex.

### Conformational destabilization of EF-Tu catalyzed by ^R^Hsp33

Structural determinants discriminating the proteolytic competence between ^Mono^EF-Tu and ^Oligo^EF-Tu were investigated by spectroscopic experiments. Notably, the far-UV circular dichroism (CD) spectrum of ^Oligo^EF-Tu, compared with that of ^Mono^EF-Tu, was characterized by a gain in random-coil and β-sheet contents at the significant expense of α-helical content (Fig 4A), which is usually described as partial unfolding and aggregation. The minute spectral change of ^Mono^EF-Tu after 30 min at 30°C (inset in Fig 4A) was also attributable to the spontaneous production of a small portion of ^Oligo^EF-Tu (see green trace in Fig 1A) that probably entailed partial unfolding. The fluorescence spectrum of ^Oligo^EF-Tu-bound 8-anilino-1-naphthalene sulfonic acid (ANS) (green trace in Fig 4B) showed a markedly higher intensity and a substantial (>25 nm) blue shift compared to the ^Mono^EF-Tu-bound ANS emission (black trace in Fig 4B), indicating a change in the tertiary structure with expansion of the hydrophobic surfaces.

We next monitored the Hsp33-induced conformational change of EF-Tu. The CD spectrum of the ^R^Hsp33:^Mono^EF-Tu complex was nearly identical to the theoretical sum of individual protein spectra (compare blue trace and open circles in Fig 4C), indicating no significant conformational change upon binding. However, subsequent incubation of the mixture resulted in a significant spectral change (red trace in Fig 4C; decreased and increased signals at around 220 nm and 200 nm, respectively) indicative of partial unfolding. The unfolding of ^Mono^EF-Tu alone occurred gradually over incubation time (black trace in the inset of Fig 4C), whereas the ^R^Hsp33:^Mono^EF-Tu complex rapidly underwent intense unfolding (red trace in the inset of Fig 4C). The ANS fluorescence spectrum of the ^R^Hsp33:^Mono^EF-Tu complex (blue trace in Fig 4B) also showed a significant blue shift and intensification after 30 min of incubation (red trace in Fig 4B). Although Hsp33 undergoes partial unfolding upon oxidation that exposes hydrophobic surfaces (Graf *et al,* 2004; Raman *et al,* 2001), such Hsp33 unfolding by unexpected oxidation was unlikely under our experimental condition, as the isolated ^R^Hsp33 showed no significant spectral change during incubation (gray trace in the inset of Fig 4C). The proteolytic resistance of ^R^Hsp33 in complex with EF-Tu (Fig 3B), in contrast to the efficient digestion of ^hO/O^Hsp33 by Lon (Fig EV4), could also exclude the possibility of Hsp33 unfolding during incubation of the ^R^Hsp33:^Mono^EF-Tu complex.

The NMR results provided more convincing evidence for the EF-Tu-specific conformational change. Although the readily precipitating property of EF-Tu at the high concentration for NMR prohibited obtaining a high-quality spectrum, the measured NMR spectrum of ^Mono^EF-Tu showed many resolved peaks that reflect a well-ordered conformation (black in Fig 4D). The ^R^Hsp33-titrated EF-Tu spectrum (red in Fig 4D) also showed overall line broadening as observed in the EF-Tu-titrated ^R^Hsp33 spectrum (Fig 2B). However, upon complex formation, chemical shift perturbations that are relevant to unfolding were evident for the remaining resonances of EF-Tu (Fig 4D), whereas ^R^Hsp33 resonances retained their chemical shifts (Fig 2B and 5A). Collectively, the spectroscopic inspection revealed that ^R^Hsp33, without its own conformational change, catalyzed the conformational change of EF-Tu to a misfolded state. In addition, given that the Lon protease preferentially recognizes hydrophobic patches in misfolded proteins (Van Melderen & Aertsen, 2009; Vieux *et al*, 2013), prominent digestion of ^R^Hsp33-bound EF-Tu (Fig 3B) could also be attributed to its aberrant folding that exposes hydrophobic surfaces (Fig 4B).

**Figure 5.**
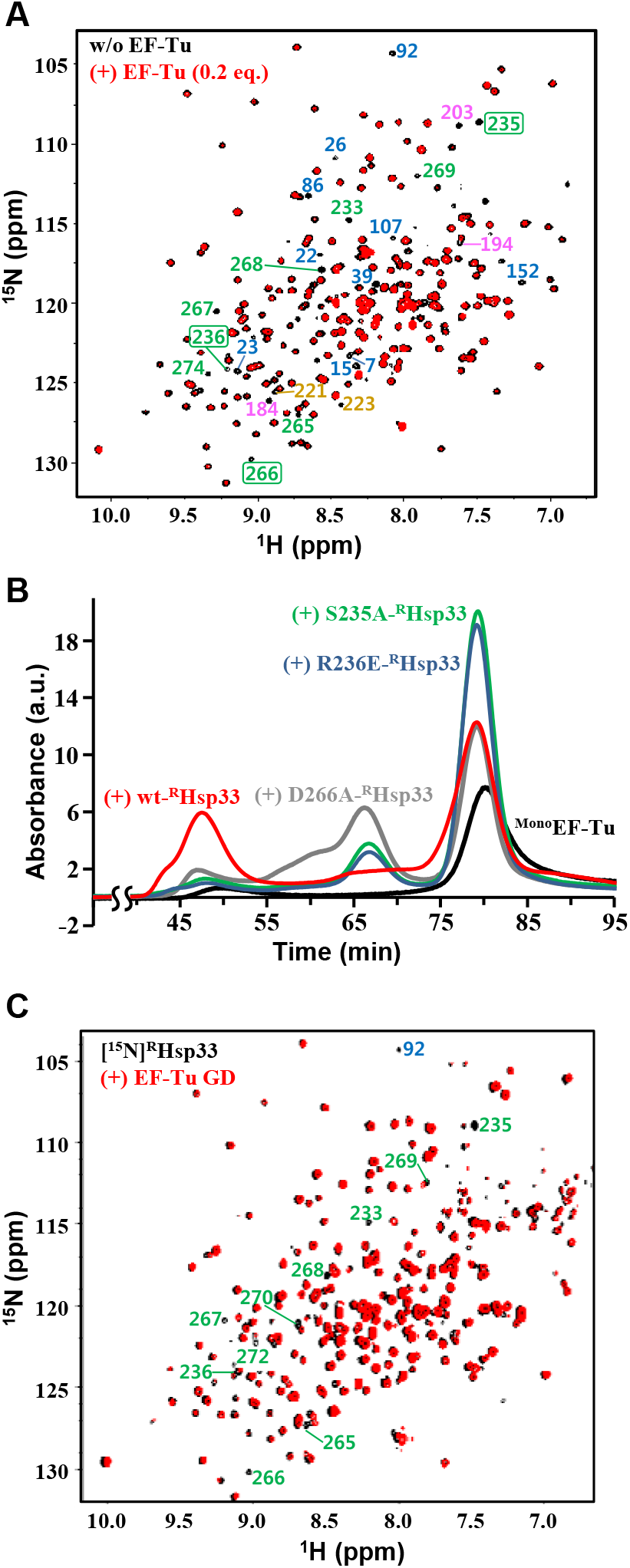
Critical contribution of the zinc-binding region in ^R^Hsp33 to EF-Tu binding. **A.** NMR ([^1^H/^15^N]TROSY) spectra of [^15^N]^R^Hsp33 (0.3 mM) at pH 6.5 in the absence (black) and presence (red) of 0.2 equimolar EF-Tu. In the well-resolved regions, indicator peaks (i.e., completely disappearing resonances upon binding) for the N-terminal core domain, MLD, interdomain linker stretch, and RSD are labeled with corresponding residue numbers colored blue, pink, tan, and green, respectively; those selected for mutagenesis are boxed. **B.** Gel-filtration (Superdex 200 column) profiles of ^Mono^EF-Tu (30 μM; black) and its mixture with equimolar wild-type (red) or a site-directed mutant of ^R^Hsp33: D266A- (gray), R236E- (blue), S235A-RHsp33 (green). All samples contained 40 μM of GDP and were pre-incubated at 30°C for 1 h before injection. **C.** NMR ([^1^H/^15^N]TROSY) spectra of [^15^N]^R^Hsp33 (0.3 mM) at pH 7.0 in the absence (black) and presence (red) of equimolar EF-Tu GD. In the well-resolved regions, unambiguous assignments are labeled for significantly affected resonances by the EF-Tu GD.

### Mutual interaction between the RSD of ^R^Hsp33 and the G-domain of EF-Tu

Owing to the immense line broadening in the NMR spectrum of the equimolar ^R^Hsp33:EF-Tu complex (Fig 2B), the EF-Tu-interacting residues of ^R^Hsp33 were qualitatively traced in the presence of a small (0.2 equimolar) amount of EF-Tu. The spectrum distinguished some representative residues of ^R^Hsp33 with their corresponding resonances disappearing upon the addition of EF-Tu (Fig 5A). Given that these residues are presumably involved in the specific molecular interaction, the EF-Tu-binding sites appeared to be distributed through all three domains of ^R^Hsp33. However, it was particularly noteworthy that many residues in the RSD, including the zinc-liganding cysteine C265, are involved in the putative EF-Tu-contacting sites, since the zinc-bound RSD fold critically discriminates ^R^Hsp33 from the EF-Tu binding-defective form ^hO/O^Hsp33. Therefore, among the putative EF-Tu-contacting residues in the RSD, we conducted site-directed mutagenesis for S235, R236, and D266, as these residues are commonly surface-exposed (Fig EV1A), are adjacent to the zinc-coordinating cysteines C234 and C265, and show a high degree of conservation in Hsp33 orthologs (Fig EV1B; serine at position 235, a positively charged amino acid at 236, and a polar amino acid at 266). Compatibility of the RSD to its zinc-bound fold was guaranteed by unaltered CD spectra of the three variants generated (Fig EV1C): S235A (hydroxyl group removed), R236E (charge reversed), and D266A (charge removed) mutants of ^R^Hsp33. Both the S235A and R236E mutations dramatically impaired the ability of ^R^Hsp33 to catalyze the EF-Tu oligomerization, while D266A also moderately corrupted the unfoldase/aggregase activity of the protein (Fig 5B). The considerable defects in aggregating EF-Tu due to directed mutagenesis at these three sites verified that the zinc-binding region of ^R^Hsp33 critically contributes to its specific binding to EF-Tu.

To identify the ^R^Hsp33 RSD-interacting domains of EF-Tu, we attempted to prepare the following five recombinant proteins corresponding to individual domains of EF-Tu and their deletion variants: GD (G-domain, residues 1–205), domain-II (D2; residues 204–294), domain-III (D3; residues 294–393), GD-deleted (ΔGD) variant (^ΔGD^EF-Tu; D2+D3), and D3-deleted (ΔD3) variant (^ΔD3^EF-Tu; GD+D2). Among them, D2 and D3 were completely insoluble, while the others that could be obtained as soluble proteins showed relatively poor stability (more rapid precipitation during storage) than intact EF-Tu. Nonetheless, the EF-Tu variant-titrating NMR analysis to monitor some obvious alterations in ^R^Hsp33 resonances was permitted at different pH values that are relatively more favorable for each EF-Tu variant. As a result, significantly broadened resonances in the EF-Tu GD-titrated ^R^Hsp33 spectrum (Fig 5C) were mapped mostly onto the RSD, including the selected mutagenesis positions (S235, R236, and D266). These resonance alterations in the RSD were also well-defined in the ^ΔD3^EF-Tu-titrated spectrum (Fig EV5A), whereas they were not relevant in the ^ΔGD^EF-Tu-titrated spectrum (Fig EV5B). Therefore, the EF-Tu GD was reasonably identified as the specific domain responsible for binding to the RSD of ^R^Hsp33. The ^ΔD3^EF-Tu (GD+D2)-titrated spectrum, which showed more abundant resonance perturbations than the GD-titrated spectrum, indicated that D2 also contributes to the ^R^Hsp33 binding of EF-Tu. In addition, the apparently weakened affinity by deleting D3 (compare Fig EV5A for ^ΔD3^EF-Tu with Fig 2B and 5A for intact EF-Tu) implied that D3 in intact EF-Tu is also possibly involved in the ^R^Hsp33 binding. Collectively, these results suggest cooperative binding of all three domains in intact EF-Tu to accomplish the observed strong binding to ^R^Hsp33 and a subsequent conformational change. However, given the severe defect in ^R^Hsp33 binding of ^ΔGD^EF-Tu (Fig EV5B), it is also reasonable that the EF-Tu GD binding to the ^R^Hsp33 RSD could drive the whole molecular interaction of the two proteins. This assumption is in turn strongly underpinned by the previous mutagenesis results (Fig 5B) that confirmed a critical influence of the mutation at the RSD of ^R^Hsp33 on its EF-Tu binding capability.

### *In vivo* relevance of EF-Tu aggregation and its effect on cell growth

To detect the ^R^Hsp33-induced aggregation of EF-Tu *in vivo*, we employed the *E. coli* JH13 strain (Cremers *et al*, 2010; Graumann *et al*, 2001) that lacks both the Lon-coding (*lon*) and Hsp33-coding (*hsl*O) genes. Cells transformed with the Hsp33-expressing plasmid (pET21a-Hsp33) were cultured under reducing conditions and the EF-Tu contents in the pellets and supernatants of the cell lysates were separately quantified and compared to the corresponding levels in total lysates. The results showed a significant increase of the pellet-fraction EF-Tu level upon expression of Hsp33, which was particularly remarkable when the cells were treated with a period of heat shock during the culture (Fig 6A). Furthermore, to evaluate whether the insoluble EF-Tu aggregates act as a noxious substance in the cell, we compared the cell growth of various *E. coli* strains after heat shock that could be readily overcome by a normal wild type *E. coli* strain (ATCC 25922) (left panel in Fig 6B). The growth of strains JH13 (Δ*lon*, Δ*hsl*O) and JH13t (JH13 transformed with pET21a-Hsp33) commonly lacking expression of both Lon and Hsp33 was similar to that of wild-type *E. coli*, which was likely overexpressing both Hsp33 and Lon during the heat-shock period. In sharp contrast, the BL21(DE3)pLysS strain showed a severe defect in cell growth; the BL21(DE3)pLysS strain was expected to overexpress Hsp33 without Lon upon heat shock, since it was used as one of the *E. coli* B strains lacking the *lon* gene. Likewise, the inducer (IPTG)-mediated overexpression of Hsp33 in the JH13t strain inhibited cell growth in a manner dependent on the inducer concentration. These results implied that overexpression of the molecular chaperone Hsp33 in the absence of Lon could be harmful to cell growth. In addition, the capability of Hsp33 to interact with EF-Tu correlated with the cellular toxicity of Hsp33 overexpression, as the variants overexpressing Hsp33 with defects in the EF-Tu interaction (i.e., a point mutation or deletion of the RSD) showed relatively weaker toxic effects (middle panel in *Figure 6B*). Considering the effect of wild-type Hsp33 to inhibit cell growth by approximately 83% (red bar versus blue bar in the right panel of *Figure 6B*), the RSD-deleted (ΔRSD) Hsp33 only caused 44% cell growth inhibition. The S235A (65% inhibition) and R236E (72% inhibition) mutations in the RSD, which significantly corrupted the ability of Hsp33 to aggregate EF-Tu (*Figure 5B*), also resulted in significantly weakened toxicity to cell growth. In contrast, the D266A-Hsp33 mutant, which showed only moderate impairment of EF-Tu-aggregating activity (*Figure 5B*), showed a comparable level (80%) of cell growth inhibition to that of wild-type Hsp33. Collectively, these results indicate that the EF-Tu-aggregating activity of Hsp33 observed *in vitro* could also be relevant *in vivo*, and its cellular effect is deleterious in the absence of Lon protease.

**Figure 6.**
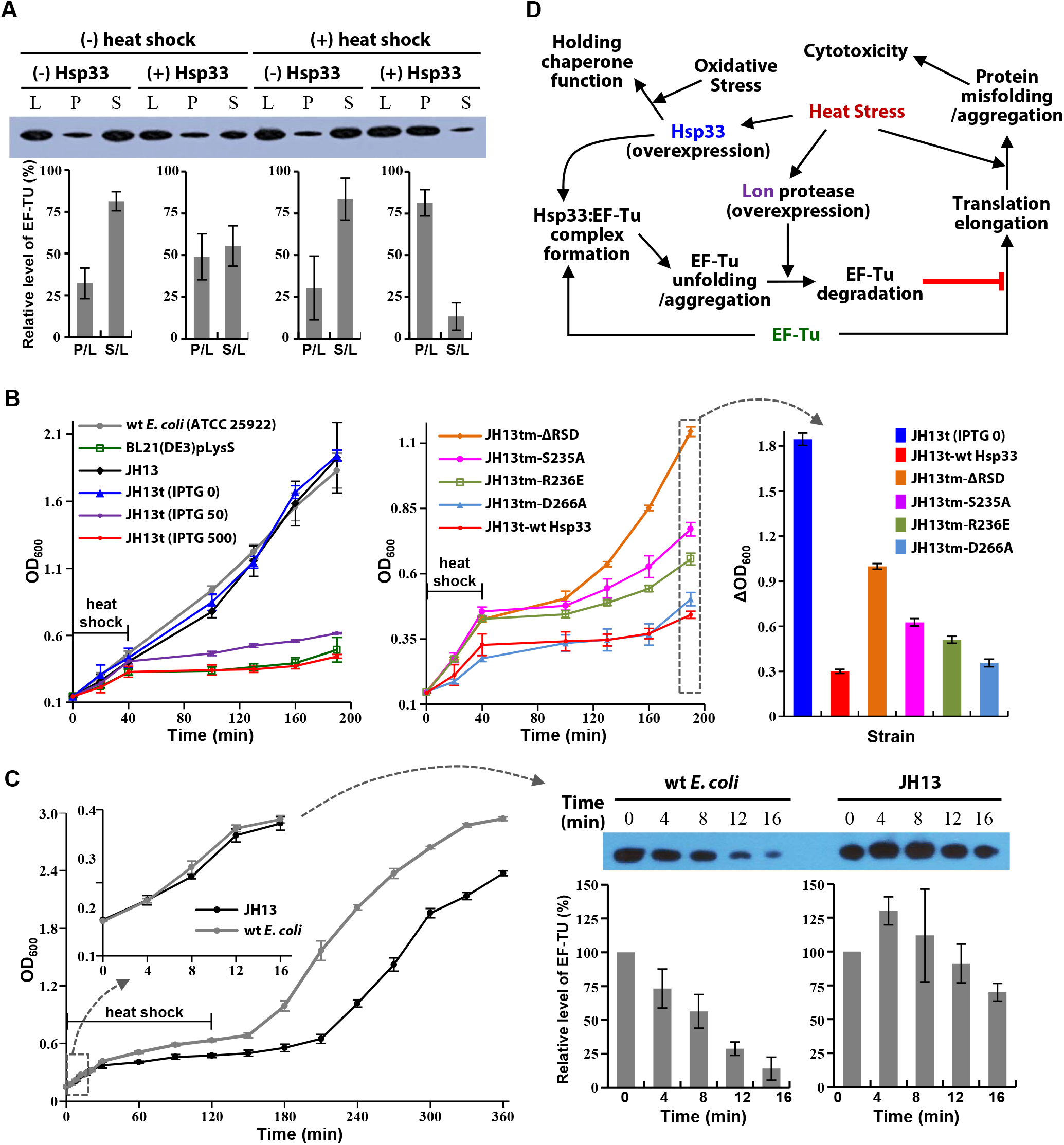
*In vivo* effects of ^R^Hsp33-induced EF-Tu aggregation. **A.** Detection and quantification of insoluble EF-Tu by immunoblotting. Levels of EF-Tu were compared between JH13t cells grown at 37°C without any stimulus and cells treated with heat shock (43°C, 40 min) and/or Hsp33 overexpression in the pellet (P) and supernatant (S), calculated relative to the level in the lysate (L). Error bars represent standard deviations of five independent experiments (n = 5). **B.** Effects of Hsp33 expression on cell growth. Growth of indicated E. coli strains during and after heat shock (43°C, 40 min) was monitored by measuring the OD600. Left panel, Hsp33 overexpression was induced by 50 or 500 μM of IPTG; middle panel, wild-type and the indicated variants of Hsp33 were overexpressed by 500 μM of IPTG; right panel, increments of OD600 (ΔOD600) during 190 min of culture. Error bars represent standard deviations (n = 3 each). **C.** Comparison of cell growth and EF-Tu levels between wild-type E. coli and the JH13 strain. Growth of the two strains during and after heat shock (45°C, 120 min) was monitored by measuring OD600 (error bars, standard deviations; n = 3 each). The growth curves at the initial stage (0–16 min) are enlarged in the inset and the normalized levels of EF-Tu at every 4 min during the period are presented by immunoblot analysis in the right panel. **D.** Suggested model for EF-Tu regulation by the collaborative action of Hsp33 and Lon in heat-stressed cells. The overexpressed Hsp33 induced by heat binds to EF-Tu and catalyzes its aberrant folding, which is recognized and digested by the heat-overexpressed Lon. Consequently, such dysregulation of EF-Tu can be beneficial for bacterial survival under the heat-stressed condition, as it attenuates the translational elongation of otherwise misfolded proteins that are deleterious to cells. If oxidative stress arises, Hsp33 can be functionally converted to a holding chaperone by the oxidat

Finally, to examine the effects of the dual deletion of Hsp33 and Lon, the cellular response to an elevated temperature (45°C) for a prolonged (2 h) duration was compared between wild-type *E. coli* and the JH13 (Δ*lon*, Δ*hsl*O) strain. The JH13 mutant showed not only more prominent retardation of cell growth during heat shock but also substantially slower recovery of cell growth after relieving the heat shock (left panel in Fig 6C). Furthermore, wild-type *E. coli* showed a significant decrease in the level of cellular EF-Tu at the initial state of heat shock, whereas such down-regulation of the EF-Tu level was not evident in JH13 (right panel in Fig 6C). The observation that dual deletion of the *lon* and *hslO* genes in JH13 resulted in weakened potential of the strain for counteracting the heat-stressed condition suggests that the supposed collaboration of Hsp33 with Lon for EF-Tu down-regulation in wild-type *E. coli* would contribute to its tolerance to heat stress.

## Discussion

This study was conducted to verify a novel, additional function of the molecular chaperone Hsp33, which is potentially associated with the regulation of EF-Tu that is engaged in ribosomal protein synthesis. Originally identified as a redox-regulated holding chaperone, the zinc-bound, reduced form of Hsp33 (^R^Hsp33) has long been regarded as a functionally inactive state that is primed for oxidation-induced activation. In addition, as cellular thermal stress can readily evoke oxidative stress, it is generally considered that overexpression of ^R^Hsp33 induced by heat represents a rapid response to the oxidative stress following heat stress (Jakob *et al*, 1999). However, the present study provides an alternative interpretation that the protein in a reduced state displays its own functionality distinguished from the oxidized form. This ^R^Hsp33-specific molecular function also appears to be distinct from the reduced form-specific action of a eukaryotic chaperone Get3 (i.e., membrane targeting of clients), whose redox-regulated molecular system closely resembles that of Hsp33 (Dahl *et al*, 2015).

The ^R^Hsp33-specific functionality could be implicit in its well-ordered, zinc-bound RSD that constitutes a novel fold of the zinc-binding domain (Won *et al*, 2004). Indeed, the present results demonstrated that the RSD mediated strong (submicromolar *K*_d_) binding of ^R^Hsp33 to EF-Tu, via interacting with the G-domain of EF-Tu. Therefore, the critical involvement of a folded RSD, which is fully unfolded upon oxidation, can explain the defective binding of oxidized Hsp33 to EF-Tu. The RSD-mediated, specific binding of ^R^Hsp33 to EF-Tu also suggests that EF-Tu would be a *bona fide* client of ^R^Hsp33. Furthermore, the strong binding of ^R^Hsp33 subsequently evoked the aberrant folding of EF-Tu, which proceeded intensely after ^R^Hsp33 binding. Therefore, the unusual thermogram of ^R^Hsp33 binding to ^Mono^EF-Tu (*Figure 2C*) is a result of the unfolding process, which is generally an endothermic reaction, following the likely exothermic binding of ^R^Hsp33. The EF-Tu unfolding that also exposed hydrophobic surfaces inevitably resulted in its aggressive oligomerization, leading to irreversible aggregation. Therefore, the molecular effect of ^R^Hsp33 binding can be regarded as efficient downregulation of both the conformational (unfolding) and colloidal (aggregation) stability of EF-Tu. Given that the intrinsic unfolding/aggregation of EF-Tu was vigorous in the absence of magnesium, the ^R^Hsp33-mediated unfolding/aggregation may entail the release of bound magnesium from the G-domain of EF-Tu. However, in cells, the magnesium-free EF-Tu, which is formed by its EF-Ts binding for GDP-GTP exchange, is stabilized by the bound EF-Ts (Maracci & Rodnina, 2016; Noble & Song, 2008). Unlike the EF-Ts interaction, which occurs through the G-domain and domain-III of EF-Tu (Thirup *et al*, 2017), G-domain/domain-II or all three domains of EF-Tu appear to cooperatively interact with ^R^Hsp33 (Fig 5C, Fig EV5). Therefore, it is inferred that EF-Tu destabilization by ^R^Hsp33, irrespective of magnesium release, might be accomplished by adverse modulation of the interdomain interaction in EF-Tu, as the interdomain communication is a critical element of intact EF-Tu stability (Šanderová *et al*, 2004).

Conclusively, given that EF-Tu showed an intrinsic tendency toward unfolding/aggregation, whereas the bound ^R^Hsp33 was not subjected to such alteration, the effect of ^R^Hsp33 on EF-Tu is defined as an unfoldase and/or aggregase activity that catalyzes the aggressive unfolding/aggregation reaction of the substrate. The ^R^Hsp33-induced EF-Tu aggregation was also relevant in cells (Fig 6A). Therefore, our study provides the first example, to our knowledge, of aggregase activity displayed by a molecular chaperone. The unfoldase activity of ^R^Hsp33, which targets the functional native fold of the specific substrate EF-Tu, is also distinctive from that of other known molecular chaperones such as GroEL and AAA+ ring proteases, which act on stable misfolded polypeptides (Mattoo *et al*, 2014; Priya *et al*, 2013; Van Melderen *et al*, 2009; Vieux *et al*, 2013). In addition, the AAA+ ring proteases, including Lon, require ATP for their unfoldase activity that is coupled with subsequent proteolytic activity (Van Melderen *et al*, 2009; Vieux *et al*, 2013), whereas the unfoldase action of Hsp33 does not consume ATP. In the case of the chaperonin GroEL, ATP-independent unfoldase activity can be exerted in connection with a subsequent ATP-consuming reaction for client refolding (Priya *et al*, 2013), whereas the unfoldase action of ^R^Hsp33 results in aggregation of the substrate. In this context, Hsp33 can be appreciated as a unique example of an ATP-independent molecular chaperone that can play a distinctive dual function as an unfoldase/aggregase (i.e., ^R^Hsp33) and as a holding chaperone (i.e., ^O^Hsp33) depending on the redox status. The unfoldase/aggregase activity of ^R^Hsp33, which is contradictory to the holding functionality of ^O^Hsp33 preventing the client aggregation by pausing the unfolding process, was achieved via its specific interaction with the natively folded EF-Tu, in contrast to the sequence non-specific, promiscuous interaction of ^O^Hsp33 with universal unfolding intermediates. Therefore, it is worth searching for the specific substrates of ^R^Hsp33 other than EF-Tu to validate the client-specific functionality of ^R^Hsp33. As a recent proteomics analysis of Hsp33 identified dozens of promising Hsp33 binding partners (Thirup *et al*, 2017), their molecular interactions with ^R^Hsp33 and the structural consequences remain to be systematically explored.

Lon protease is an essential component of the proteostasis network recognizing and degrading misfolded proteins (Van Melderen *et al*, 2009; Vieux *et al*, 2013). Given that EF-Tu is a *bona fide* substrate of Lon protease in cells (Bruel *et al,* 2012), the unfoldase/aggregase action of ^R^Hsp33 on EF-Tu clearly functions to make EF-Tu recognizable by Lon, as the natively folded EF-Tu was not susceptible to proteolytic degradation by Lon (Fig 3A). Therefore, in the absence of Lon, the expression of ^R^Hsp33 results in the cellular accumulation of insoluble EF-Tu (Fig 6A), which in turn adversely affects cell growth (Fig 6B). The collaborative action of ^R^Hsp33 and Lon to degrade EF-Tu can also be harmful to cell growth unless prevented and/or compensated by another regulatory system. Indeed, Bruel et al. observed that the Hsp33 overproduction in a DnaK-deficient *E. coli* strain exhibited strong toxicity to bacterial growth under normal conditions by up-regulating the Lon-mediated degradation of EF-Tu (Bruel *et al*, 2012). Conversely, however, growth of the DnaK-deficient *E. coli* at non-permissive temperatures was rescued by overexpressing Hsp33 (Bruel *et al*, 2012), which implies that the EF-Tu degradation by ^R^Hsp33 and Lon could be beneficial for bacterial survival under stressful conditions. Finally, we detected substantial down-regulation of EF-Tu in wild-type *E. coli* upon heat shock, which was not observed in the JH13 mutant (Δ*lon*, Δ*hsl*O) strain. Moreover, dual deletion of Hsp33- and Lon-encoding genes (JH13) resulted in a phenotype that was less tolerant to the severe heat stress that wild-type *E. coli* is able to surmount (Fig 6C). Therefore, given that both Hsp33 and Lon belong to the heat shock regulon of bacteria (Van Melderen *et al*, 2009; Jakob *et al,* 1999), we suggest a mechanism for the plausible contribution of ^R^Hsp33 to cell survival in heat shock, which is schematically presented in Fig 6D. This model also considered the fact that global pausing of elongation is a widespread cellular translation regulation mechanism for cell survival in heat shock (Shalgi *et al*, 2013), since heat stress promotes the noxious misfolding and aggregation of proteins synthesized in ribosome. In the suggested model (Fig 6D), the Hsp33-bound EF-Tu would certainly lose its functionality for translation elongation as a result of the aberrant conformational change. Subsequently, the EF-Tu-specific degradation by Lon would enable Hsp33 to be recycled to further attenuate the translation elongation. Therefore, given the overexpression of ^R^Hsp33 and Lon by heat shock, the present results demonstrating the ^R^Hsp33-catalyzed degradation of EF-Tu by Lon may implicate ^R^Hsp33 in the elongation-posing machinery for cell survival in response to thermal stressors, which can also contribute to proteostasis by regulating the proteome-wide turnover rate in cells.

This possible *in vivo* regulation of EF-Tu by ^R^Hsp33 would be linked to other molecular systems, particularly those including the foldase chaperone DnaK. Crosstalk with DnaK is also required for the redox-regulated holding function of Hsp33 to turn its bound substrates (i.e., the oxidation-induced unfolding intermediates) over to DnaK when ^O^Hsp33 reverts to ^R^Hsp33 upon subsided oxidative stress (Reichmann *et al*, 2012). However, reversed crosstalk would operate for the proposed action of ^R^Hsp33 in heat shock, since EF-Tu is a likely client of DnaK under normal conditions; e.g., the eukaryotic counterpart of DnaK (Hsp70) has been verified to interact with eukaryotic EF-Tu (EEF1A1) under normal conditions and to act as a central repressor of elongation pausing (Shalgi *et al*, 2013). Given that the interaction between the cognate pair of EF-Tu-DnaK (EEF1A1-Hsp70) in eukaryotes was confirmed to be significantly weakened during heat stress (Shalgi *et al*, 2013), our results showing the heat-sensitive triggering of EF-Tu along with the strong binding of ^R^Hsp33 to the unfolded EF-Tu (Fig 2C) suggest that heat shock could serve as an effective signal for EF-Tu to switch its binding chaperone from DnaK to ^R^Hsp33. In addition to this plausible competition between DnaK and ^R^Hsp33 for client binding, trigger factor (TF) has also been suggested to be involved in the chaperone network (Bruel *et al*, 2012), probably by facilitating the Hsp33-mediated EF-Tu degradation by Lon; however, it is not yet established whether ^R^Hsp33, TF, or any other chaperones in cells can deliver EF-Tu to Lon. Furthermore, the ribosome-dissociated EF-Tu in cells is normally stabilized by EF-Ts for its guanine nucleotide exchange (Maracci & Rodnina, 2016; Noble & Song, 2008; Thirup *et al*, 2015). Therefore, to gain systematic insight into the EF-Tu regulon in cells, the interplay mode of these related chaperones should be further investigated at the structural and molecular interaction levels. In addition, as mentioned above, the recently proposed Hsp33 interactome comprises dozens of non-stress induced proteins, the majority of which are involved in protein biogenesis pathways (Rimon *et al*, 2017). Therefore, the specific cellular functionality of ^R^Hsp33 is expected to be increasingly revealed through further studies on the individual molecular interactions of ^R^Hsp33 with these putative clients. Moreover, considering the fact that Hsp33 is expressed at a basal level even under non-stressed conditions (Dahl *et al*, 2015), the ^R^Hsp33-specific functionality might also be engaged as a component of the cellular PQC machinery under normal conditions. However, it remains to be verified whether the EF-Tu regulation by ^R^Hsp33 is compatible with such housekeeping roles.

EF-Tu in *Cyanobacterium synechocystis* has been identified as a target of oxidation by cellular reactive oxygen species, resulting in oligomerization/aggregation (Yutthanasirikul *et al*, 2016). Considering that Hsp33 unfolds upon oxidation for its holding activity, a functional switch of Hsp33 for prevention of EF-Tu aggregation under oxidative stress might be possible (Fig 6D), which can address the intriguing observation that *E. coli* Hsp33 could protect against EF-Tu degradation in *Vibrio cholerae* under oxidative conditions (Bruel *et al*, 2012). However, Lon is unlikely to be involved in this currently uncharacterized system, since the oxidized Hsp33 could also be an efficient substrate of the protease (Fig EV4). Alternatively, the chaperone network, including Hsp33 and Lon, for the PQC machinery might be differently modulated depending on the organism and/or the type of cellular stress, which also remains to be investigated.

In summary, this study highlighted an unfoldase activity of the molecular chaperone Hsp33 that catalyzes the structural conversion of EF-Tu to an aggregation- and proteolysis-prone state. Therefore, the present results provide the first evidence of aggrease activity or unique ATP-independent unfoldase activity displayed by a molecular chaperone to realize its substrate aggregation. In addition, the intriguing kinetics (Fig 4C) and thermodynamics (Fig 2C) of EF-Tu unfolding/aggregation upon ^R^Hsp33 binding suggest that this paradigm could serve as an excellent model system for further in-depth analyses of the protein unfolding pathway. With respect to functional aspects of Hsp33, the holding-inactive, reduced conformation of Hsp33 was specifically relevant to the interaction and destabilization of EF-Tu, which otherwise is unlikely targeted to rapid digestion by its cognate quality-control machinery, Lon protease. This novel molecular functionality, also collaborating with Lon protease, is expected to be linked to the translational elongation pause occurring during heat shock, thereby regulating protein turnover for proteostasis in cells to allow them to endure the heat-stressed conditions. The functional significance of the EF-Tu dysregulation by Hsp33 is worthy of further investigation, considering the multifunctional properties of EF-Tu (Yutthanasirikul *et al*, 2016), including its chaperone activity, beyond its known involvement in protein biosynthesis.

## Material and Methods

### DNA constructs

All primer sequences used for subcloning and mutagenesis are summarized in Table S1. DNA fragments encoding *E. coli* Hsp33 were PCR-amplified from the plasmid pUJ30 (Jakob *et al*, 1999) as a template and inserted into the pET21a vector between the *Nde*I and *Xho*I restriction sites. The reverse primer contained a stop codon to produce the protein without artificial histidine tags. For the *E. coli* EF-Tu construct, the corresponding open reading frame was PCR-amplified using the genomic DNA of the *E. coli* BL21(DE3)pLysS strain as a template, followed by insertion into the pCold-I (Takara) vector between the *Nde*I and *Xho*I restriction sites, which conferred an N-terminal hexahistidine tag on the expressing protein. These constructed plasmids for wild-type Hsp33 and EF-Tu were used as templates for subsequent point mutations of Hsp33, which were introduced using the Phusion Site-Directed Mutagenesis Kit (Finnzymes), and subcloning of EF-Tu variants, respectively. Following verification by DNA sequencing, the recombinant plasmids for Hsp33 mutants were transformed into the *E. coli* strain JH13 (BL21, Δ*hsl*O) (Graumann *et al*, 2001), whereas *E. coli* BL21(DE3)pLysS cells transformed with the constructs for EF-Tu and its individual variants were used for their expression, respectively.

### Protein preparation

For preparation of Hsp33 and its mutants, the transformed cells were grown in Luria–Bertani (LB) medium at 37°C until the optical density at 600 nm (OD600) reached about 0.7, followed by induction of the expression by adding isopropyl-β-D-1-thiogalactopyranoside (IPTG) and ZnSO4, both at a final concentration of 1 mM. After prolonged (6 h) induction, the cells were harvested and resuspended in ice-cold standard buffer (50 mM Tris-HCl, 50 mM NaCl, pH 7.4) containing 5 mM dithiothreitol (DTT) and 50 μM ZnSO4. The cells were disrupted by sonication, and the proteins were purified from the supernatant via the sequential application of anion-exchange, hydrophobic-interaction, and gel-permeation chromatography. During the purification, 5 mM DTT and 50 μM ZnSO4 were continuously contained in the standard buffer to ensure maintenance of the expressed proteins as reduced, zinc-bound forms. For oxidation, the purified protein solution was buffer-exchanged by passing through a PD-10 column (GE Healthcare) to remove the residual DTT and zinc ions. Oxidation of the purified protein (50 μM) was then performed by incubating at 43°C for 3 h in the presence of H2O2 (2 mM) as an oxidant. Individual species of oxidized forms were separately pooled from the elution through gel filtration (Lee *et al*, 2012; Lee *et al*, 2015). Each protein solution was buffer-exchanged with an appropriate buffer for subsequent experiments by diafiltration using Amicon Ultra-15 devices (Merck Millipore). The protein concentration was determined spectrophotometrically using molar absorptivity at 280 nm deduced from the amino acid sequence of each protein.

For EF-Tu and its variants, the protein expression was induced with IPTG (1 mM) and MgSO4 (1 mM) at 17°C for 18 h. Cell lysis buffer contained 50 mM Tris-HCl (pH 7.4), 1 mM DTT, 10 mM MgSO4, 70 mM imidazole, and 100 mM NaCl. Protein purification was performed via the sequential application of Ni^2+^-affinity, anion-exchange, and gel-permeation chromatography in the standard buffer containing 1 mM DTT and 5 mM MgSO4. For rapid oligomerization of EF-Tu, residual DTT and magnesium ions were removed from the purified ^Mono^EF-Tu solution by using the PD-10 column (GE Healthcare), followed by treatment of the solution with 1 mM EDTA. ^Oligo^EF-Tu was then collected from the supernatant of the EDTA-treated solution for subsequent spectroscopic analysis. The other procedures followed the protocols described above for Hsp33.

### Analytical gel filtration

Gel filtration was performed on a HiLoad 16/600 Superdex^™^ 75 or Superdex^™^ 200 column (GE Healthcare) connected to an fast protein liquid chromatography (FPLC) system at a flow rate of 1 mL/min in 50 mM HEPES-OH buffer (pH 7.4) containing 150 mM NaCl, 5 mM MgSO4, 100 μM ZnSO4, and 5 mM DTT. The injection volume of each analyte was approximately 2 mL and the eluting proteins were detected by measuring absorbance at 280 nm. The hydrodynamic size of each protein was represented by the apparent molecular mass (kDa), which was deduced from the elution volume, in comparison with molecular mass standards of the gel filtration calibration kits (GE Healthcare), LMW (for Superdex 75) and HMW (for Superdex 200).

### Pull-down assay

Thirty microliters of Ni^2+^-charged His-Bind Agarose Resin suspension (ELPisBio, Korea) pre-equilibrated in a standard buffer (50 mM HEPES-OH, pH 7.4, 50 mM NaCl, 5 mM MgSO4, 100 μM ZnSO4, 1.3 mM β-mercaptoethanol) was mixed with the bait protein (hexahistidine-tagged EF-Tu) solution in the same buffer to a final volume of 100 μL and a final bait concentration of 30 μM, followed by 4°C incubation for 15 min. After three washes (repeated adding buffer and spin down of resin) with the standard buffer containing 70 mM imidazole, 100 μL of prey solution (60 μM ^R^Hsp33, 80 μM ^hO^Hsp33, or 80 μM ^O^Hsp33) was added to the bait-bound resin, followed by end-over-end mixing and subsequent 1-h incubation at 4°C. The resin suspension was then washed three times with 100 μL of standard buffer containing 70 mM imidazole and 0.4% (v/v) NP-40. Bound materials were then eluted with 100 μL of standard buffer containing 600 mM imidazole. After spin-down removal of the resin, the supernatant (eluted solution) was resolved by sodium dodecyl sulfate-polyacrylamide gel electrophoresis (SDS-PAGE).

### Lon-proteolysis assay

One hundred microliters of substrate protein (EF-Tu and/or Hsp33) solution (10 μM) in 20 mM sodium phosphate buffer (pH 7.4) containing 50 mM NaCl, 100 μM MgSO4, 50 μM ZnSO4, 100 μM ATP, and 5 μM pyruvate kinase (Merck) was reacted with 0.26 nM of recombinant *E. coli* Lon protease (Sino Biological) at 30°C. To halt the Lon reaction, 1 μL of EDTA stock (100 mM) and 5 μL of Protease Inhibitor Cocktail (Merck) stock (1 mg/mL) solution were added into every sample (10 μL) taken at designated sampling times, followed by boiling with SDS-PAGE sample buffer (10 μL) and subsequent storage in a deep freezer until use for the SDS-PAGE run.

### Light scattering analysis

The ^Oligo^EF-Tu (100 μM) solution in 50 mM HEPES-OH buffer (pH 7.4) containing 150 mM NaCl and 5 mM DTT was thoroughly filtered using a membrane filter (Advantec) with a pore diameter of 0.2 μm. Dynamic light scattering data of the solution was collected on a Viscotek 802 DLS instrument (Malvern Instruments), followed by the molecular weight determination using OmniSIZE software (Malvern Instruments). The aggregation of ^Mono^EF-Tu (100 μM) at 30°C in the absence and presence of half-equimolar ^R^Hsp33 was monitored by recording the kinetic traces of light scattering from the protein solution at 400 nm (5-nm slit width for both excitation and emission), using a Varian Cary Eclipse spectrofluorophotometer with continuous stirring. The solvent buffer (pH 7.4) contained 50 mM HEPES-OH, 50 mM NaCl, 5 mM MgSO4, 100 μM ZnSO4, and 5 mM DTT.

### ITC analysis

Binding thermodynamics were measured at 25°C using a MicroCal Auto-iTC200 calorimeter. EF-Tu (70 μM) solutions in 50 mM HEPES-OH buffer (pH 7.4) containing 50 mM NaCl, 5 mM MgSO4, 50 μM ZnSO4, and 5 mM DTT were contained in the reaction cell (200 μL), whereas Hsp33 (210 μM) solutions in the same buffer were titrated from the syringe (40 μL). A titration experiment consisted of 20 injections: 0.4 μL of the first injection followed by 19 injections (2 μL each), with an injection interval of 150 s. The obtained thermogram was analyzed by data fitting on a single-site binding model, using a commercial software package (ORIGIN 7.0) provided by the manufacturer.

### CD spectroscopy

A 0.1-cm path length cell was used for the CD measurements of individual protein samples (5–20 μM) dissolved in 15 mM sodium phosphate buffer (pH 7.4) containing 15 mM NaCl, 20 μM ZnSO4, and 20 μM MgSO4. Standard far-UV CD spectra were recorded on a Jasco J-710 spectropolarimeter at room temperature (approximately 22°C) with a 1-nm bandwidth and a 1-s response time. Three individual scans taken from 260 nm to 190 nm with a 0.1- or a 1-nm step resolution were summed and averaged, followed by subtraction of blank buffer CD signals. Time-course CD changes were monitored at 220 nm at 30°C, using an Applied Photophysics Chirascan CD spectrometer equipped with a temperature controller. The signals were recorded at every 0.1 s with a 1-nm bandwidth.

### Fluorescence spectroscopy

Protein solutions (5 μM) containing no or 11 μM of the fluorescent probe ANS (Merck) were prepared in 50 mM HEPES-OH buffer (pH 7.4) containing 50 mM NaCl, 5 mM DTT, 5 mM MgSO4, and 100 μM ZnSO4. Fluorescence spectra were recorded in a Varian Cary Eclipse spectrofluorophotometer at 30°C with continuous stirring. The excitation wavelength was fixed at 370 nm (slit width 1 nm), while fluorescence emissions were scanned from 400 to 600 nm (slit width 1 nm).

### NMR spectroscopy

The isotope-[^15^N]-enriched proteins for NMR measurements were produced by growing the expressing cells in M9 minimal medium supplemented with ^15^NH4Cl as the sole nitrogen source. NMR samples contained 0.3 mM of the [^15^N]-labeled target protein and varying concentrations of its interacting non-labeled counterpart, dissolved in 50 mM HEPES-OH buffer (pH 7.4) containing 50 mM NaCl, 5 mM MgSO4, 1 mM ZnSO4, 5 mM DTT, and 7% (v/v) D2O. Conventional [^1^H/^15^N]-TROSY spectra were measured at 298 K on a Bruker Biospin Avance 900 spectrometer equipped with a cryoprobe. The previously assigned chemical shift values (Lee *et al*, 2015) were used for residue-specific inspection of the ^R^Hsp33 spectra.

### Cell growth assay and immunoblotting of EF-Tu

To detect the ^R^Hsp33-induced aggregation of EF-Tu, various strains of *E. coli* cells were grown at 37°C in 70 mL of LB broth medium containing 6 mM of L-cysteine as a reducing agent (Ohtsu *et al*. 2010). When the OD600 reached around 0.15, the cells were treated with heat shock at 43°C for 40 min, along with induction of Hsp33 by adding 50 or 500 μM of IPTG. Subsequent to the heat shock, the cells were further incubated at 37°C for 2 h, followed by sampling (5 mL each) for immunoblotting. The sampled cells harvested by centrifugation were lysed in PBS (75 μL per 0.1 of OD600) by sonication. Pellets and supernatants from half of the cell lysates were separated by centrifugation. The pellets were washed with PBS three times, followed by re-suspension in PBS at the same volume of the supernatant sample. Finally, 2% of the prepared lysate, pellet, and supernatant samples were subjected to SDS-PAGE. Resolved proteins on the gel were then transferred onto a polyvinylidene difluoride membrane (Millipore), followed by overnight incubation at 4°C with anti-EF-Tu antibodies (1:50 dilution; HycultBiotech) in 5% bovine serum albumin containing 0.05% Tween-20 and subsequent incubation with horseradish peroxidase-conjugated secondary antibodies (1:500) for 1 h at room temperature. The membrane was then developed using an ECL Prime kit (Translab) and exposed on CL-G PLUS medical X-ray film (Agafa HealthCare NV). The relative intensities of the protein bands were quantified by densitometric analysis using ImageJ software. Tolerance to heat shock was compared between wild-type *E. coli* (ATCC 25922 strain) and the JH13 strain by cultivation at 45°C for 120 min, followed by prolonged incubation at 37°C. Cell lysates of the samples taken every 4 min for the initial 16 min of heat-shock cultivation were subjected to the immunoblot analysis of EF-Tu in the same manner described above.

## Acknowledgements

This work was supported by the National Research Foundation (NRF) (grant no. 2010-0006022, 2013R1A1A2007774, and 2016R1A2B4009700) funded by the Korean government (MEST). The use of NMR, CD, fluorescence, and ITC equipment was supported by the Korea Basic Science Institute (Ochang, Korea) under the R&D program (Project No. D37700) supervised by the Ministry of Science and ICT. We thank Dr. U. Jakob (University of Michigan, USA) for generously providing the recombinant plasmid pUJ30 and the *E. coli* strain JH13.

## Author contribution

K.S.J., J.S.K, C.Y.W, and Y.S.L constructed the recombinant plasmids and prepared protein samples under the supervision of M.D.S and H.S.W. K.S.J., J.H.K, and K.S.R. performed the NMR experiments and data analysis. K.S.J. and Y.H.L performed ITC experiments and data analysis. K.S.J. and Y.S.L performed biochemical and other spectroscopic experiments. J.H.K., K.S.R., M.D.S., Y.H.L., and H.S.W. conducted the data analysis. H.S.W. conceived of the study and directed the experiments and data analysis. K.S.J. and H.S.W wrote the paper.

## Conflict of interests

The authors declare that no competing interests exist.

---

**Figure EV1:**
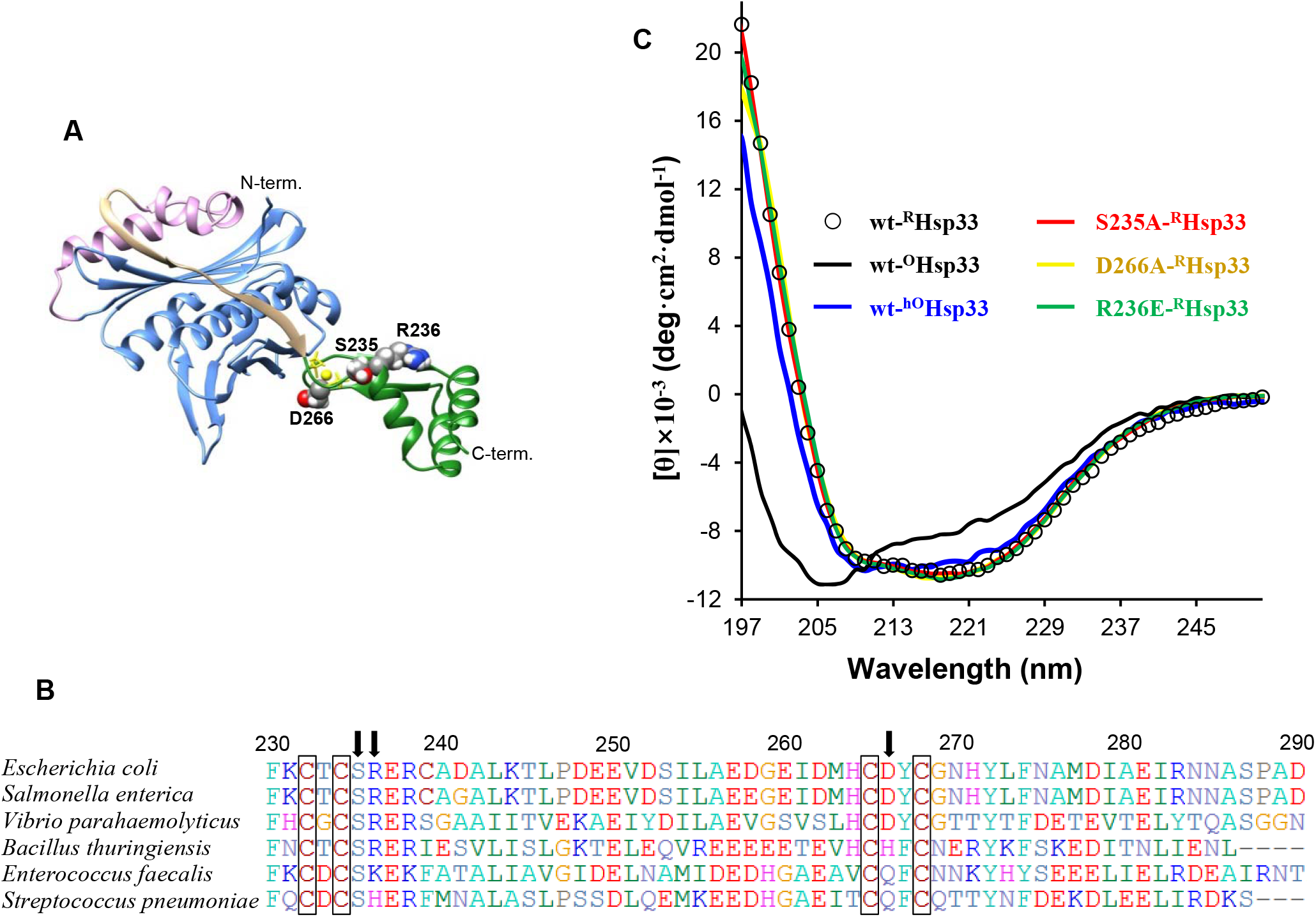
Selected positions for the site-directed mutagenesis of Hsp33. (**A**) Semi-empirical model structure of *E. coli* ^R^Hsp33 determined previously (*22*). The NCD, MLD, interdomain linker stretch, and RSD are colored blue, pink, tan, and green, respectively. The bound zinc ion (sphere) and the conserved Cys sidechains (stick models) are depicted in yellow. Sidechains of the S235, R236, and D266 residues for mutagenesis are represented as spheres. (**B**) Amino acid conservations in the RCDs of selected Hsp33 homologs. Multiple sequence alignments were performed with CLUSTALW (www.genome.jp/tools-bin/clustalw) and edited with the BioEdit program (www.mbio.ncsu.edu/BioEdit/bioedit.html). Zinc-coordinating cysteines are boxed and the selected positions for mutagenesis are indicated by arrows. Sequence numbers are for *E. coli* Hsp33. (**C**) Maintained conformation of ^R^Hsp33 mutants. Normalized ([θ], mean residue molar ellipticity) far-UV CD spectra of D266A- (yellow), S235A- (red), and R236E-^R^Hsp33 (green) are superimposed. The previously reported CD spectra of wild-type Hsp33^20^ are represented for comparison: ^R^Hsp33, open circles; ^hO^Hsp33, blue line; ^O^Hsp33, black line.

**Figure EV2:**
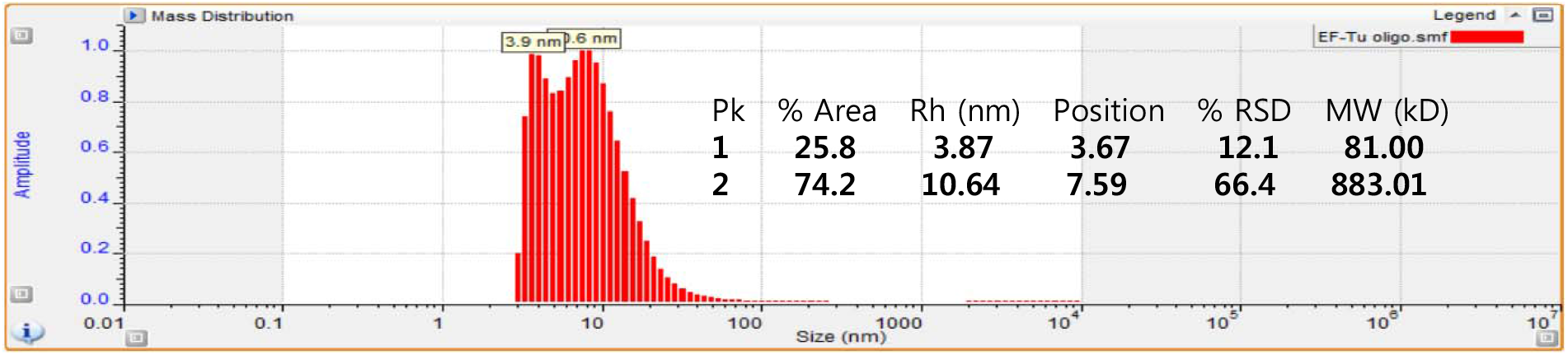
DLS monitoring of EF-Tu oligomers. Oligomeric fraction of purified EF-Tu prepared inthe absence of Mg^2+^ was measured before the ITC experiments (see Figure 2C). Analyzed parameters are presented as an inset table.

**Figure EV3:**
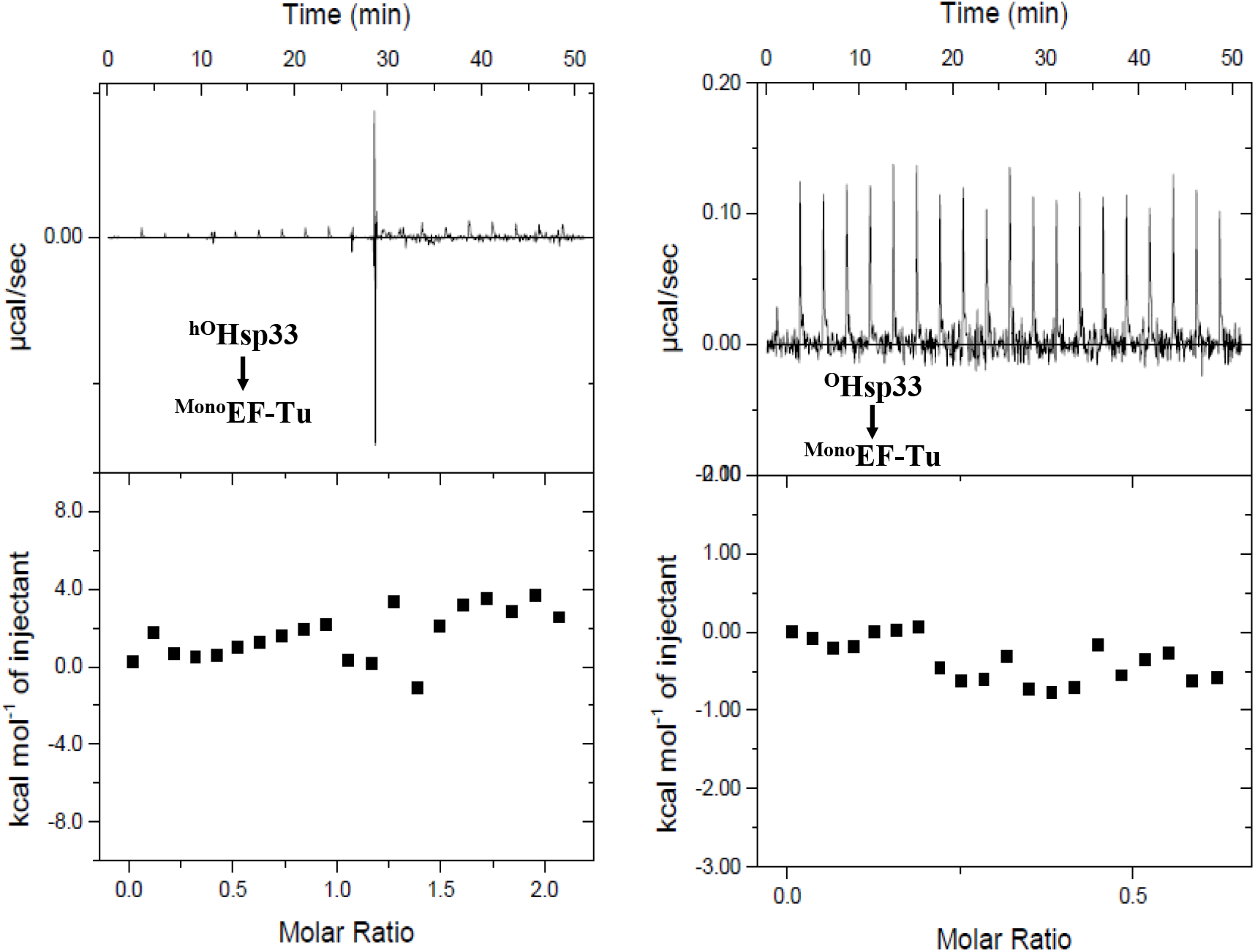
ITC measurements for ^hO^Hsp33 (left) and ^O^Hsp33 (right) titration to ^Mono^EF-Tu. Each point in the binding isotherm (lower panels) represents the integrated heat of the associated peak in the thermogram (upper panels).

**Figure EV4:**
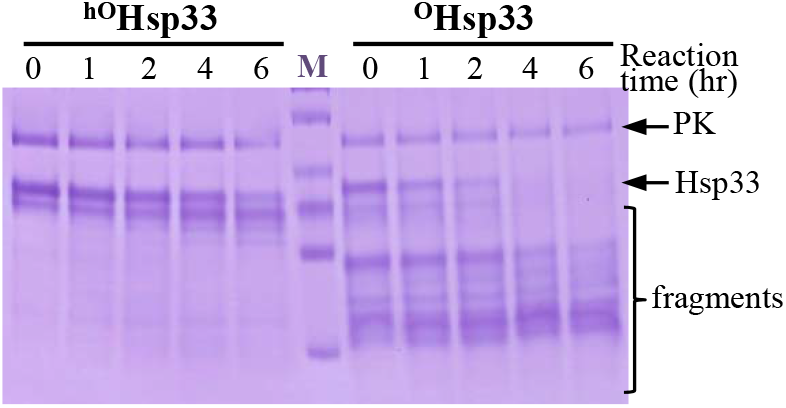
Digestion of oxidized Hsp33 by Lon protease. ^hO^Hsp33 and ^O^Hsp33 (10 μM each) were reacted at 30°C with 0.26 nM Lon protease supplemented with pyruvate kinase (PK; 5 μM) for ATP generation. The 0-hr sample was taken immediately after adding the protease into the reaction mixture.

**Figure EV5:**
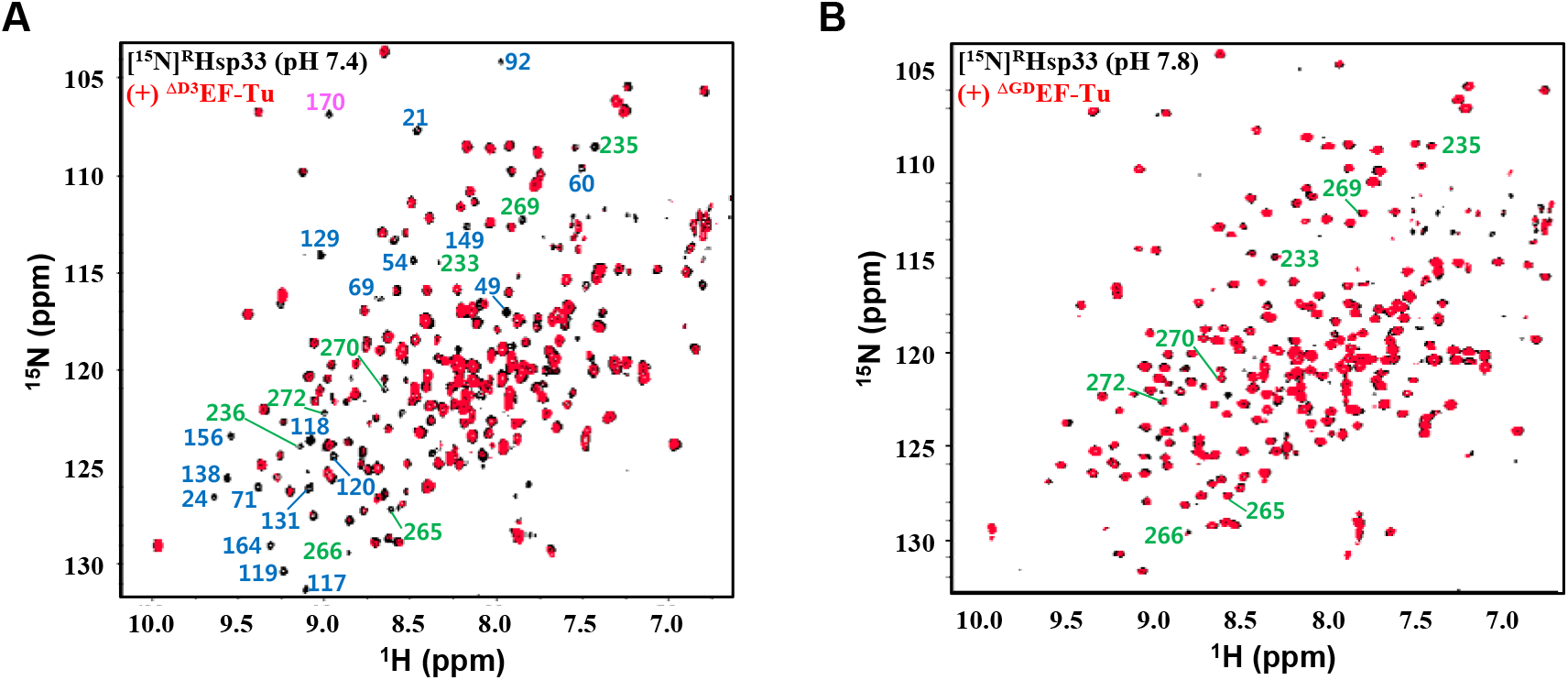
NMR ([^1^H/^15^N]TROSY) spectra of [^15^N]^R^Hsp33 (0.3 mM) in the absence (black) and presence (red) of equimolar ^ΔD3^EF-Tu (A; pH 7.4) and 2 equimolar ^ΔGD^EF-Tu (B; pH 7.8). In the well-resolved regions, unambiguous assignments are labeled for significantly affected resonances by the ^ΔD3^EF-Tu (**A**) or for selected indicator peaks from RSD (**B**). Coloring designation follows that described in Figure 5A.

**Table S1.**
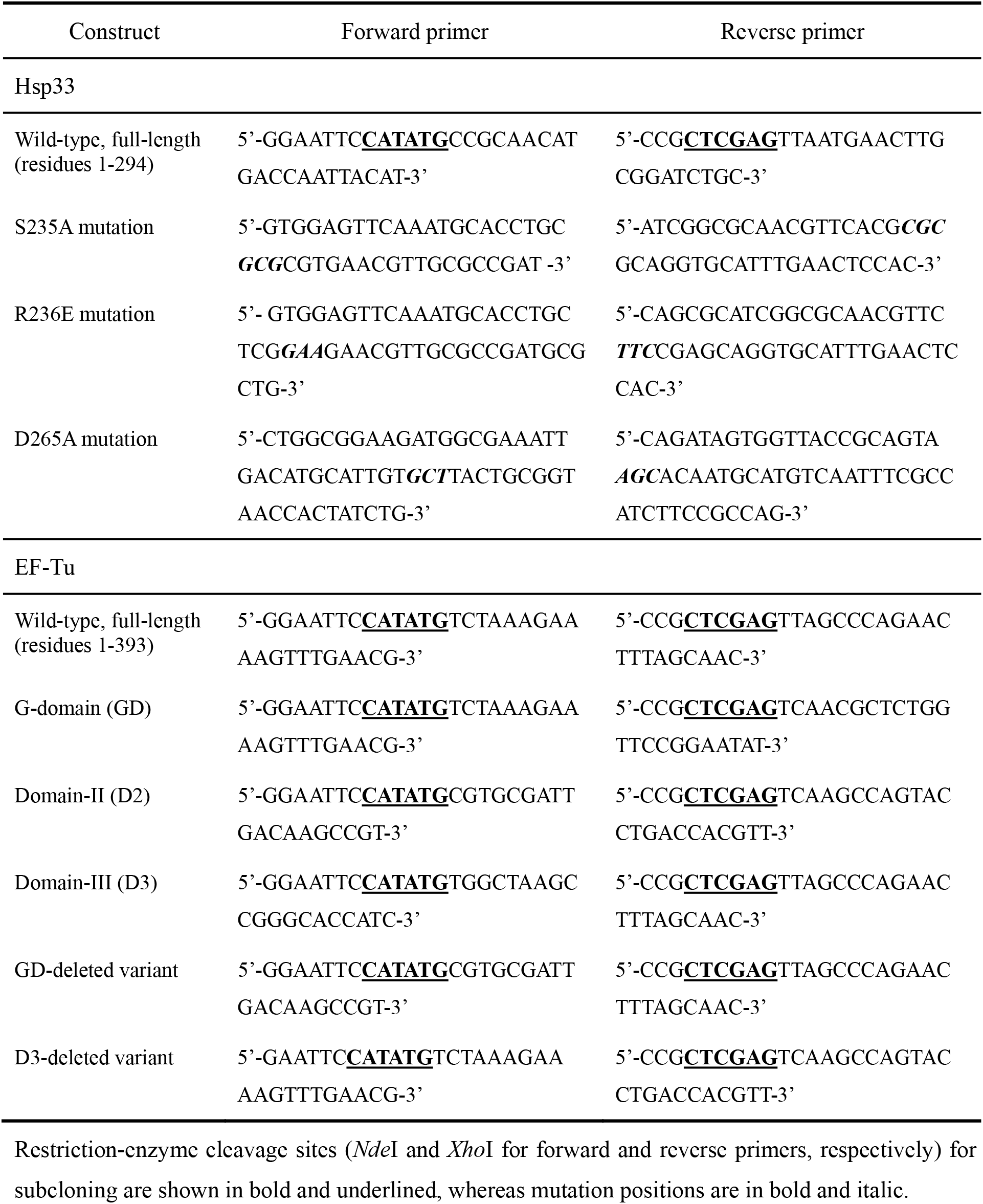
Primer sequences employed for subcloning and mutagenesis.

## References

Balchin D, Hayer-Hartl M, Hartl FU (2016) In vivo aspects of protein folding and quality control. Science 353: Aac4354.

Bershtein S, Mu W, Serohijos AWR, Zhou J, Shakhnovich EI (2013) Protein quality control acts on folding intermediates to shape the effects of mutaions on organismal fitness. Mol Cell 49: 133–144

Bruel N, Castanié-Cornet MP, Cirinesi AM, Koningstein G, Georgopoulos C, Luirink J, Genevaux P (2012) Hsp33 controls elongation factor-Tu stability and allows *Escherichia coli* growth in the absence of the major DnaK and trigger factor chaperones. J Biol Chem 287: 44435–44446

Cremers CM, Reichmann D, Hausmann J, Ilbert M, Jakob U (2010) Unfolding of metastable linker region is at the core of Hsp33 activation as a redox-regulated chaperone. J Biol Chem 285: 11243–11251

Craig EA, Weussman JS, Horwich AL (1994) Heat shock proteins and molecular chaperones:mediators of protein conformation and turnover in the cell. Cell 78: 365–372

Doyle SM, Genest O, Wickner S (2013) Protein rescue from aggregates by powerful molecular chaperone. Nat Rev Mol Cell Biol 14: 617–629

Dahl JU, Gray MJ, Jakob U (2015) Protein quality control under oxidative stress conditions. J Mol Biol 427, 1549–1563

Fu J, Momcilovic I, Prasad PVV (2012) Roles of protein synthesis elongation factor EF-Tu in heat tolerance in plants. J Botany 835836

Graumann J, Lilie H, Tang X, Tucker KA, Hoffmann JH, Vijayalakshmi J, Saper M, Bardwell JCA, Jakob U (2001) Activation of the redox-regulated molecular chaperone Hsp33-a two-step mechanism. Structure 9: 377–387

Graf PCF, Martinez-Yamout M, VanHaerents S, Lilie H, Dyson HJ, Jakob U (2004) Activation of the redox-regulated chaperone Hsp33 by domain unfolding. J Biol Chem 279: 20529–20538

Groitl B, Horowitz S, Makepeace KA, Petrotchenko EV, Borchers CH, Reichmann D, Bardwell JC, Jakob U (2016) Protein unfolding as a switch from self-recognition to high-affinity client binding. Nat Commun 7: 10357

Hartl FU, Bracher A, Hayer-Hartl M (2011) Molecular chaperones in protein folding and proteostasis. Nature 475: 324–332

Ilbert M, Horst J, Ahrens S, Winter J, Graf PCF, Lilie H, Jakob U (2007) The redox-switch domain of Hsp33 functions as dual stress sensor. Nat Struct Mol Biol 14: 556–563

Jakob U, Muse W, Eser M, Bardwell JCA (1999) Chaperone activity with a redox switch. Cell 96: 341–352

Lee YS, Lee J, Ryu KS, Lee Y, Jung TG, Jang JH, Sim DW, Kim EH, Seo MD, Lee KW, Won HS (2015) Semi-empirical structure determination of *Escherichia coli* Hsp33 and identification of dynamic regulatory elements for the activation process. J Mol Biol 427: 3850–3861

Klaips CL, Jayaraj GG, Hartl FU (2018) Pathways of cellular proteostasis in aging and disease. J Cell Biol 217: 51–63

Lee YS, Ryu KS, Kim SJ, Ko HS, Sim DW, Jeon YH, Kim EH, Choi WS, Won HS (2012) Verification of the interdomain contact site in the inactive monomer, and the domain-swapped fold in the active dimer of Hsp33 in solution. FEBS Lett 586: 411–415

Maracci C, Rodnina MV (2016) Translational GTPases. Biopolymers 105: 463–475

Mattoo RUH, Goloubinoff P (2013) Molecular chaperones are nanomachines that catalyticallyt unfold misfolded and alternatively folded proteins. Cell Mol Life Sci 71: 3311–3325

Noble CG, Song H (2008) Structural studies of elongation and release factors. Cell Mol Life Sci 65: 1335–1346

Ohtsu I, Wiriyathanawudhiwong N, Morigasaki S, Nakatani T, Kadokura H, Takagi H (2010) The L-cysteine/L-systine shuttle system provides reducing equivalents to the periplasm in *Escherichia coli*. J Biol chem 285: 17479–17487

Priya S, Sharma SK, Sood V, Mattoo RU, Finka A, Azem A, De Los Rios P, Goloubinoff P (2013) GroEL and CCT are catalytic unfoldases mediating out-of-cage polypeptide refolding without ATP. Proc Natl Acad Sci USA 110: 7199–7204

Priya S, Sharma P, Goloubinoff P (2013) Molecular chaperones as enzymes that catalytically unfold misfolded polypeptides. FEBS Lett 587: 1981–1987

Raman B, Kumar LVS, Ramakrishna T, Rao CM (2001) Redox-regulated chaperone function and conformational changes of *Escherichia coli* Hsp33. FEBS Lett 489: 19–24

Reichmann D, Xu Y, Cremers CM, Ilbert M, Mittelman R, Fitzgerald MC, Jakob U (2012) Order out of disorder: working cycle of an intrinsically unfolded chaperone. Cell 148: 947–957

Rimon S, Suss O, Goldenberg M, Fassler R, Yogev O, Amartely H, Propper G, Friedler A, Reichmann D (2017) A role of metastable regions and their connectivity in the inactivation of a redox-regulated chaperone and its inter-chaperone crosstalk. Antioxid Redox Signal 27: 1252–1267

Šanderová H, Hùlková M, Maloñ P, Kepková M, Jonák J (2004) Thermostability of multidomain proteins: elongation factors EF-Tu from *Escherichia coli* and *Bacillus stearothermophilus* and their chimeric forms. Protein Sci 13: 89–99

Shalgi R, Hurt JA, Krykbaeva I, Taipale M, Lindquist S, Burge CB (2013) Widespread regulation of translation by elongation pausing in heat shock. Mol Cell 49: 439–452

Taipale M, Tucker G, Peng J, Krykbaeva I, Lin ZY, Larsen B, Choi H, Berger B, Gingras AC, Lindquist S (2014) A quantitative chaperone interaction network reveals the architecture of cellular protein homeostasis pathways. Cell 158: 434–448

Thirup SS, Van LB, Nielsen TK, Knudsen CR (2015) Structural outline of the detailed mechanism for elongation factor Ts-mediated guanine nucleotide exchange on elongation factor Tu. J Struct Biol 191: 10–21

Van Melderen L, Aertsen A (2009) Regulation and quality control by Lon-dependent proteolysis. Res Microbiol 160: 645–651

Vieux EF, Wohlever ML, Chen JZ, Sauer RT, Baker TA (2013) Distinct quaternary structures of the AAA+ Lon protease control substrate degradation. Proc Natl Acad Sci USA 110: E2002–E2008

Visscher M, De Henau S, Wildschut MHE, van Es RM, Dhondt I, Michels H, Kemmeren P, Nollen EA, Braeckman BP, Burgering BMT, Vos HR, Dansen TB (2016) Proteome-wide changes in protein turnover rates in *C. elegans* models of longevity and age-related disease. Cell Rep 16: 3041–3051

Voth W, Jakob U (2017) Stress-activated chaperones: a first line of defense. Trends Biochem Sci 42: 899–913

Wholey WY, Jakob U (2012) Hsp33 confers bleach resistance by protecting elongation factor Tu against oxidative degradation in *Vibro cholerae*. Mol Microbiol 8: 981–991

Winter J, Ilbert M, Graf PC, Ozcelik D, Jakob U (2008) Bleach activates a redox-regulated chaperone by oxidative protein unfolding. Cell 135: 691–701

Won HS, Low LY, De Guzman R, Martinez-Yamout M, Jakob U, Dyson HJ (2004) The zinc-dependent redox switch domain of the chaperone Hsp33 has a novel fold. J Mol Biol 341: 893–899

Yutthanasirikul R, Nagano T, Jimbo H, Hihara Y, Kanamori T, Ueda T, Haruyama T, Konno H, Yoshida K, Hisabori T, Nishiyama Y (2016) Oxidation of a cysteine residue in elongation factor EF-Tu reversibly inhibits translation in the cyanobacterium *Synechocystis* sp. PCC 6803. J Biol Chem 291: 5860–5870

